# SLC33A1 exports oxidized glutathione to maintain endoplasmic reticulum redox homeostasis

**DOI:** 10.64898/2026.02.01.703113

**Authors:** Shanshan Liu, Mark Gad, Caifan Li, Kevin Cho, Yuyang Liu, Khando Wangdu, Viktor Belay, Alon Millet, Hiroyuki Kojima, Henry Sanford, Michele Wölk, Linas Urnavicius, Maria Fedorova, Gary J. Patti, Ekaterina V. Vinogradova, Richard K. Hite, Kıvanç Birsoy

**Affiliations:** Laboratory of Metabolic Regulation and Genetics, The Rockefeller University, New York, NY, USA; Structural Biology Program, Memorial Sloan Kettering Cancer Center, New York, USA; Department of Chemistry, Washington University, St Louis, MO, USA; Center for Mass Spectrometry and Metabolic Tracing, Washington University, St Louis, MO, USA; Siteman Cancer Center, Washington University School of Medicine, St Louis, MO, USA; Center for Human Nutrition, Department of Medicine, Washington University School of Medicine, St Louis, MO, USA; Physiology, Biophysics, and Systems Biology (PBSB) Program, Weill Cornell Graduate School of Biomedical Sciences, New York, NY, USA; Computational and Systems Biology Program, Memorial Sloan Kettering Cancer Center, New York, NY, USA; Laboratory of Systems Cancer Biology, The Rockefeller University, New York, NY, USA; Laboratory of Chemical Immunology and Proteomics, The Rockefeller University, New York, NY, USA; Center of Membrane Biochemistry and Lipid Research, University Hospital and Faculty of Medicine Carl Gustav Carus of Dresden University of Technology, Dresden, Germany; Laboratory of Chemistry and Cell Biology, The Rockefeller University, New York, NY, USA

## Abstract

The endoplasmic reticulum (ER) requires an oxidative environment to support the efficient maturation of secretory and membrane proteins. This is in part established by glutathione, a redox-active metabolite present in reduced (GSH) and oxidized (GSSG) forms. The ER maintains a higher GSSG:GSH ratio than the cytosol; however, the mechanisms controlling ER redox balance remain poorly understood. To address this, we developed a method for the rapid immunopurification of the ER, enabling comprehensive profiling of its proteome and metabolome. Combining this approach with CRISPR screening, we identified SLC33A1 as the major ER GSSG exporter in mammalian cells. Loss of SLC33A1 leads to GSSG accumulation in the ER and a liposome-based assay demonstrates that SLC33A1 directly transports GSSG. Cryo-EM structures and molecular dynamics simulations reveal how SLC33A1 binds GSSG and identify residues critical for its transport. Finally, an imbalance in GSSG:GSH ratio induces ER stress and dependency on the ER-associated degradation (ERAD) pathway, driven by a shift in protein disulfide isomerases (PDIs) toward their oxidized forms. Altogether, our work establishes SLC33A1-mediated GSSG export as a key mechanism for ER redox homeostasis and protein maturation.

## Introduction

First identified by Albert Claude in 1945^1^, the endoplasmic reticulum (ER) was later defined by George Palade through electron microscopy as a highly specialized membrane-bound organelle essential for the synthesis, folding, and maturation of secretory and membrane proteins^2^. The ER also functions as a metabolic hub, supporting N-glycosylation, calcium storage, and the biosynthesis of phospholipids, cholesterol, and sphingolipids^3–5^. To sustain these reactions and ensure high-quality protein output, cells tightly control the ER’s metabolic and chemical environment^6^. However, the full repertoire of metabolites and enzymatic reactions within the ER remains incompletely understood. Among these, redox-active metabolites are critical for balancing the oxidative and reductive processes required during protein folding and preventing the accumulation of misfolded proteins^7^. Disruptions in this balance trigger ER stress responses that, if unresolved, contribute to diseases such as neurodegeneration and metabolic dysfunction^8^.

Glutathione, the most abundant small-molecule thiol, serves as the primary redox buffer within the ER lumen. Although synthesized in the cytosol by the consecutive actions of γ-glutamate-cysteine ligase (GCL) and glutathione synthetase (GS), glutathione is also present in the ER, which contains a small but functionally significant fraction of the total cellular pool^9^. Glutathione exists in reduced (GSH) and oxidized (GSSG) forms, with most cellular compartments maintaining a highly reducing GSSG:GSH ratio ranging from 1:10 to 1:100. By contrast, the ER sustains a more oxidative environment, with a GSSG:GSH ratio ranging from 1:1 to 1:7^9–11^, favoring disulfide bond formation. In the cytosol and mitochondria, this balance is typically established through the action of glutathione reductases, NADPH-dependent enzymes that regenerate GSH from GSSG. Notably, the ER lacks glutathione reductase activity^12^ and instead appears to rely on the transport of excess GSSG to maintain its optimal redox environment^10^. These observations suggest the existence of specialized transporters that mediate glutathione flux across the ER membrane. Despite the fundamental role of ER redox regulation in physiology and disease, the identity of ER glutathione transporters remains poorly defined.

## Results

### A rapid purification method for profiling ER metabolome and proteome

To reliably quantify ER redox metabolites, we first sought to develop a method for rapid profiling of metabolites and proteins in intact ER stacks. Because traditional centrifugation-based methods are too slow to preserve ER metabolite content, we adapted an immunopurification strategy previously developed for other organelles^13–15^. Among ER-resident transmembrane proteins screened (Extended Data Fig. 1a), we prioritized EMC3, which enabled the purification of ER with markedly reduced contamination from other organelles (Extended Data Fig. 1b). We engineered an EMC3 fusion protein comprising the fluorescent protein mScarlet and three tandem HA epitopes at its cytoplasm-facing C-terminus (EMC3-mScarlet-3xHA, ER-tag) (Fig. 1a). Immunofluorescence confirmed that the ER-tag colocalizes with ER marker calnexin (Fig. 1a). Notably, expression of ER-tag did not cause any visible changes in ER morphology (Fig. 1a) or impact ER stress response upon expression (Extended Data Fig. 1c). Immunoblotting of ER-tag immunoprecipitates (ER-IP) showed robust enrichment of ER proteins with minimal contamination from other compartments (Fig. 1b, c). Additionally, electron micrographs of ER-IP samples showed that the bead-bound fraction contains intact ER ministacks with ribosomes on the surface, verifying the specificity of the approach (Fig. 1d). Proteomic analysis further demonstrated that ER-IP selectively enriches annotated ER proteins relative to those from other compartments (Fig. 1e, f; Extended Data Fig. 1d-h). To benchmark coverage, we curated a comprehensive list of 1672 ER-associated proteins based on GO annotations as well as recently published studies^16,17^. Of ∼800 detected proteins significantly enriched over control-IPs, 508 matched entries in this curated list (Supplementary Table 1-2). Since the ER, unlike other organelles, is the primary site for the translation of secretory and membrane proteins, we also analyzed RNA content. RNAseq analysis of the ER-IP samples revealed strong enrichment (adjusted P value < 1E-10) for transcripts encoding proteins targeted to the plasma membrane, ER, Golgi, lysosome and extracellular proteins (Fig. 1g, h; Extended Data Fig. 1i, j; Supplementary Table 3).

**Fig. 1:**
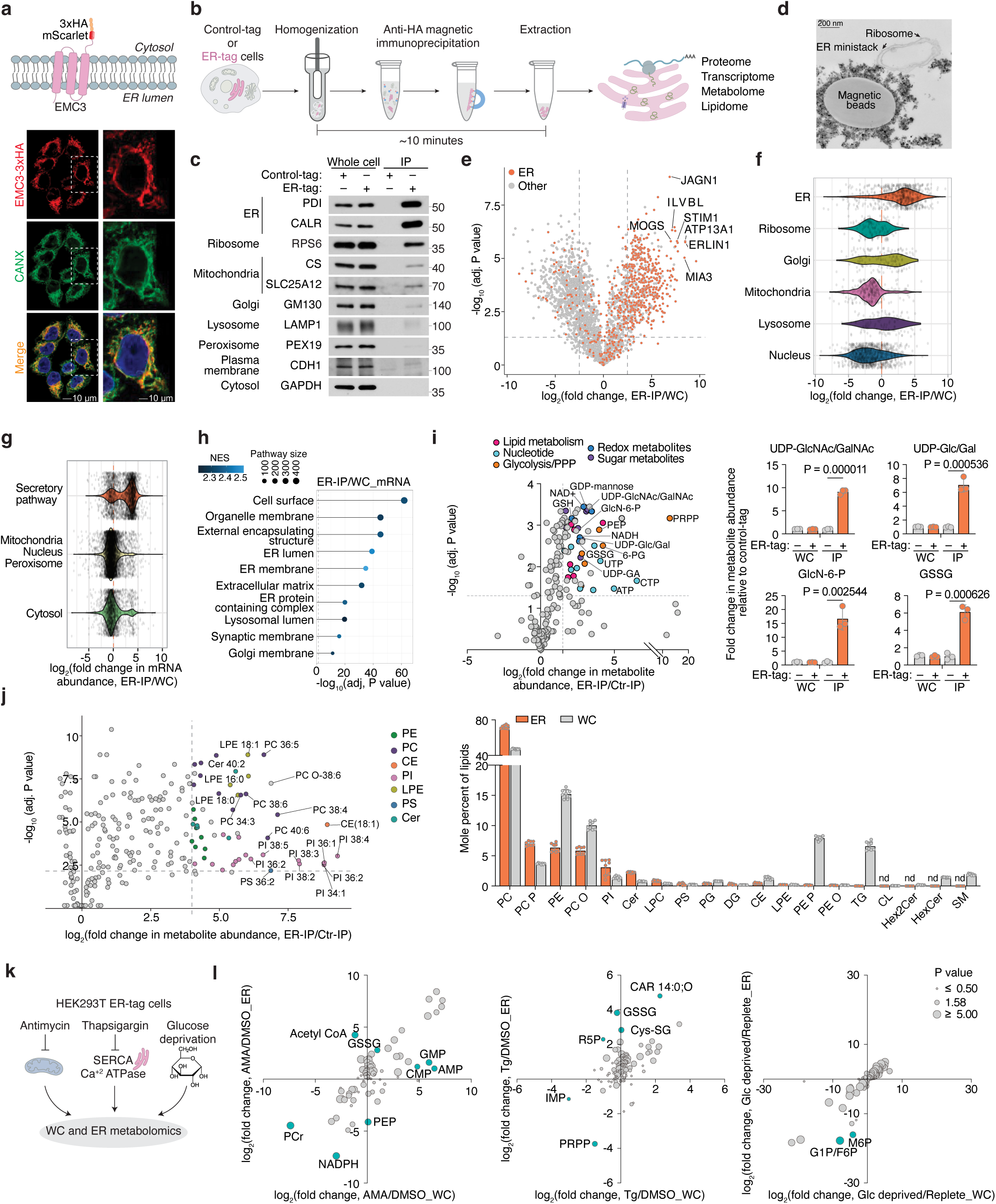
A rapid purification method for profiling ER metabolome and proteome. a. Top, schematic of the domain structure of the ER transmembrane protein EMC3 fused to mScarlet and 3×HA tag (ER-tag). Bottom, immunofluorescence analysis of ER-tag (HA, red) and calnexin (CANX, green) in HEK293T cells. Micrographs are representative of three independent experiments. Source images are available in Source Data. b. Schematic depicting the workflow for the ER-IP method. Cells expressing control-tag or ER-tag are rapidly harvested and homogenized. ER is isolated with a 5-minute IP by HA magnetic beads, washed, and then lysed for immunoblotting, or extracted for proteomic, transcriptomic, metabolomic, lipidomic analysis. c. Immunoblot analysis of whole-cell lysates (5 µg) and ER-IP lysates (5 µg) with the indicated compartment-specific antibodies. Lysates were derived from HEK293T cells expressing control-tag or ER-tag. Immunoblots are representative of three independent experiments with similar results. Unprocessed blots are available in Source Data. d. Representative electron micrograph image of the immunoprecipitated ER from HEK293T cells expressing ER-tag. The intact ER ministack and ribosomes are indicated. The images are representative of three independent experiments with similar results. Source images are available in Source Data. e. Volcano plot showing relative fold change (log_2_) in protein abundance versus −log (adjusted P values) from ER-tag HEK293T cells comparing ER-IP samples to whole cell (WC) samples (n = 3). Statistical significance was assessed using a two-sided t-test with permutation-based false discovery rate (FDR) correction (1%). The dotted line represents *P*=0.05, log_2_(fold change) = 2.5. Selected highly enriched ER proteins are denoted with gene names. Source numerical data are available in Source Data. f. Violin plots showing relative fold change (log_2_) in protein abundance associated with indicated organelles from ER-tag HEK293T cells comparing ER-IP samples to whole cell samples. Source numerical data are available in Source Data. g. Violin plots showing relative fold change (log_2_) in mRNA abundance associated with indicated organelles from ER-tag HEK293T cells comparing ER-IP samples to whole cell samples. The mRNAs encoding proteins associated with secretory pathway are enriched, including ER, Golgi, lysosome, plasma membrane and extracellular matrix. Source numerical data are available in Source Data. h. Gene set enrichment analysis of significantly enriched mRNAs encoding proteins associated with secretory pathway from ER-tag HEK293T cells comparing ER-IP samples to whole cell samples. NES represents normalized enrichment score. Differential expression analysis was performed using limma-voom, and significance was determined using two-sided statistical testing with multiple-comparison correction (adjusted P value). Genes included in the analysis passed a log2(fold change) cutoff of 0.5 and adjusted P < 0.05. n = 3 biological replicates. Source numerical data are available in Source Data. i. Left, volcano plot showing relative fold change (log_2_) in metabolite abundance versus −log (adjusted P values) of ER-IP samples from ER-tag HEK293T cells compared to those from control-tag HEK293T cells. Statistical significance was determined by multiple two-tailed unpaired t-tests with correction for multiple comparisons using the Holm–Šídák method. The dotted line represents *P*=0.05, log_2_(fold change) = 1.5. UDP-GlcNAc/GalNAc, UDP-N-acetylglucosamine/ N-acetylgalactosamine; 6-PG, 6-phosphogluconic acid; GlcN-6-P, Glucosamine 6-phosphate; UDP-Glc/Gal, UDP-glucose/galactose; PEP, phosphoenolpyruvate; UDP-GA, UDP-glucuronic acid. PRPP, Phosphoribosylpyrophosphate. Right, relative metabolite abundance of indicated whole cell or ER metabolites from ER-tag HEK293T and control-tag HEK293T cells. Data are presented mean ± SD. Statistical significance was determined using a two-tailed unpaired t-test. n = 3 biological replicates. Source numerical data are available in Source Data. j. Left, volcano plot showing relative fold change (log_2_) in lipid abundance versus −log (adjusted P values) of ER-IP samples from ER-tag HEK293T cells compared to those from control-tag HEK293T cells. Statistical significance was determined by multiple two-tailed unpaired t-tests with correction for multiple comparisons using the Holm–Šídák method. The dotted line represents *P*=0.01, log_2_(fold change) = 4. Right, mole percent of indicated whole cell or ER lipids from ER-tag HEK293T cells. Data are presented mean ± SD. n = 8 biological replicates. Source numerical data are available in Source Data. k. Schematic depicting an ER metabolomics experiments using three stress conditions. l. Volcano plot showing relative fold change (log_2_) in metabolite abundance from ER-tag HEK293T under indicated stress conditions compared to untreated. The x axis shows the fold change in whole cell metabolites, and the y axis shows the fold change in ER metabolites. Statistical significance was determined by multiple two-tailed unpaired t-tests. The size of the dots represents the P value. Metabolites that change significantly differently between whole cell and ER-IP are highlighted. Antimycin: 10 µM, 2 hours, Thapsigargin: 3 µM, 4 hours. Glucose deprivation: 16 hours. n = 3 biological replicates. Source numerical data are available in Source Data.

To profile ER-specific metabolites, we used LC-MS to quantify the relative abundance of targeted polar small molecules and lipids in ER versus control immunoprecipitants. Among the detected metabolites, 53 were enriched at least two-fold in isolated ER samples. UDP-sugars, such as UDP-glucuronic acid, UDP-N-glucose/galactose, and UDP-N-acetylglucosamine (UDP-GlcNAc), which are critical for ER-associated glycosylation^18^, were strongly enriched (Fig. 1i), confirming the specificity of our metabolomics approach. Precursors for UDP-GlcNAc synthesis, including glucosamine-6-phosphate, acetyl-CoA, N-acetylglucosamine-1/6-phosphate and UTP, were similarly enriched (Fig. 1i), indicating their import into the ER or a potential activity of the hexosamine pathway in the ER. In addition, we observed metabolites from glycolysis and the pentose phosphate pathway (PPP), including glucose-6-phosphate (G6P), 6-phosphogluconate (6PG), ribose-5-phosphate, xylulose-5-phosphate, and sedoheptulose-7-phosphate (Fig. 1i; Extended Data Fig. 2a). This is consistent with studies showing that G6P is transported into the ER lumen and hydrolyzed to glucose during gluconeogenesis^19,20^, and that H6PD oxidizes G6P to 6PG, initiating the PPP and producing NADPH^21^. Redox-active metabolites, such as GSH/GSSG, NAD^+^, and NADP^+^, along with FAD, a key cofactor for ERO1 and disulfide bond formation, were also highly enriched in ER-IP samples (Fig. 1i; Extended Data Fig. 2a). Finally, we detected several nucleotides, including ATP, which supports chaperone-mediated protein folding, as well as phospholipid precursors, such as phosphoethanolamine and phosphocholine (Fig. 1i; Extended Data Fig. 2a), reflecting the ER’s central role in phospholipid synthesis. Complementary lipidomics revealed a lipid composition dominated by phosphatidylcholines (PCs), phosphatidylethanolamines (PEs), and phosphatidylinositols (PIs), in line with prior findings^5^. In contrast, sphingomyelin, glycosphingolipids, and cardiolipins were depleted, consistent with their synthesis in the Golgi apparatus and mitochondria^22^ (Fig. 1j; Extended Data Fig. 2b). Collectively, these results define the distinct biochemical landscape of the ER and demonstrate the utility of our approach for comprehensive ER metabolomics and lipidomics.

Given the compartmentalized nature of the ER, stress conditions may uniquely influence the composition of its luminal metabolites. Therefore, using the ER-IP approach, we next investigated how ER metabolites respond to distinct metabolic stressors. We profiled ER metabolites in cells treated with a mitochondrial electron transport chain inhibitor antimycin, a sarco/ER Ca^2+^-ATPase inhibitor thapsigargin (to deplete ER calcium), or subjected to glucose depletion (Fig. 1k). Antimycin selectively increased ER acetyl-CoA levels without altering total cellular pools (Fig. 1l; Extended Data Fig. 2c, d), suggesting enhanced fatty acid elongation and desaturation coupled to NAD⁺ regeneration^23^ and omega oxidation. Thapsigargin elevated ER-specific GSSG and cysteine-glutathione disulfide (Cys-SG) levels (Fig. 1l; Extended Data Fig. 2e, f), linking calcium depletion to luminal redox imbalance. Glucose withdrawal uniformly reduced glycolytic and pentose phosphate pathway intermediates in both the ER and cytosolic compartments (Fig. 1l; Extended Data Fig. 2g, h), indicating a robust exchange and metabolic connectivity between them. Collectively, the ER-IP platform enables comprehensive profiling of ER proteins, RNAs, metabolites, and lipids, facilitating the study of ER metabolism under physiological and stress conditions.

### Identification of SLC33A1 as a regulator of ER glutathione homeostasis

Given our ability to determine ER metabolites, we next sought to identify regulators of ER glutathione homeostasis in mammalian cells. We hypothesized that increasing GSH abundance specifically within the ER would create a stress state (GSH overload) and allow us to identify dependencies on potential glutathione transporters. To test this, we targeted the bacterial enzyme GshF, which combines glutamate-cysteine ligase and glutathione synthetase activities^24^, to the ER using a doxycycline-inducible system (ER-GshF) (Fig. 2a). Ectopically expressed ER-GshF in HEK293T cells efficiently localized to the ER (Fig. 2b) and produced GSH inside ER, even in the presence of buthionine sulfoximine (BSO), an inhibitor of cytosolic glutathione synthesis (Fig. 2c). ER-GshF expression did not impair cell proliferation (Extended Data Fig. 3a), suggesting the presence of compensatory mechanisms that mitigate ER GSH accumulation. To uncover these pathways, we designed an ER-focused sgRNA library and performed CRISPR-Cas9 screens in ER-GshF-expressing HEK293T cells (Fig. 2d). *SLC33A1*, an ER-localized transporter previously proposed to transport acetyl-CoA^25^, emerged as the top hit. The screens also identified ER-associated degradation (ERAD) components, including *SYVN1* and *SEL1L* (Fig. 2e; Extended Data Fig. 3b, c), implicating ERAD in the response to ER GSH overload. To validate the screen results, we generated *SLC33A1* knockout cells and expressed ER-GshF. We found increased ER GSH levels were selectively lethal in *SLC33A1* deficient cells (Fig. 2f), establishing a requirement for SLC33A1 under conditions of ER GSH overload.

**Fig. 2:**
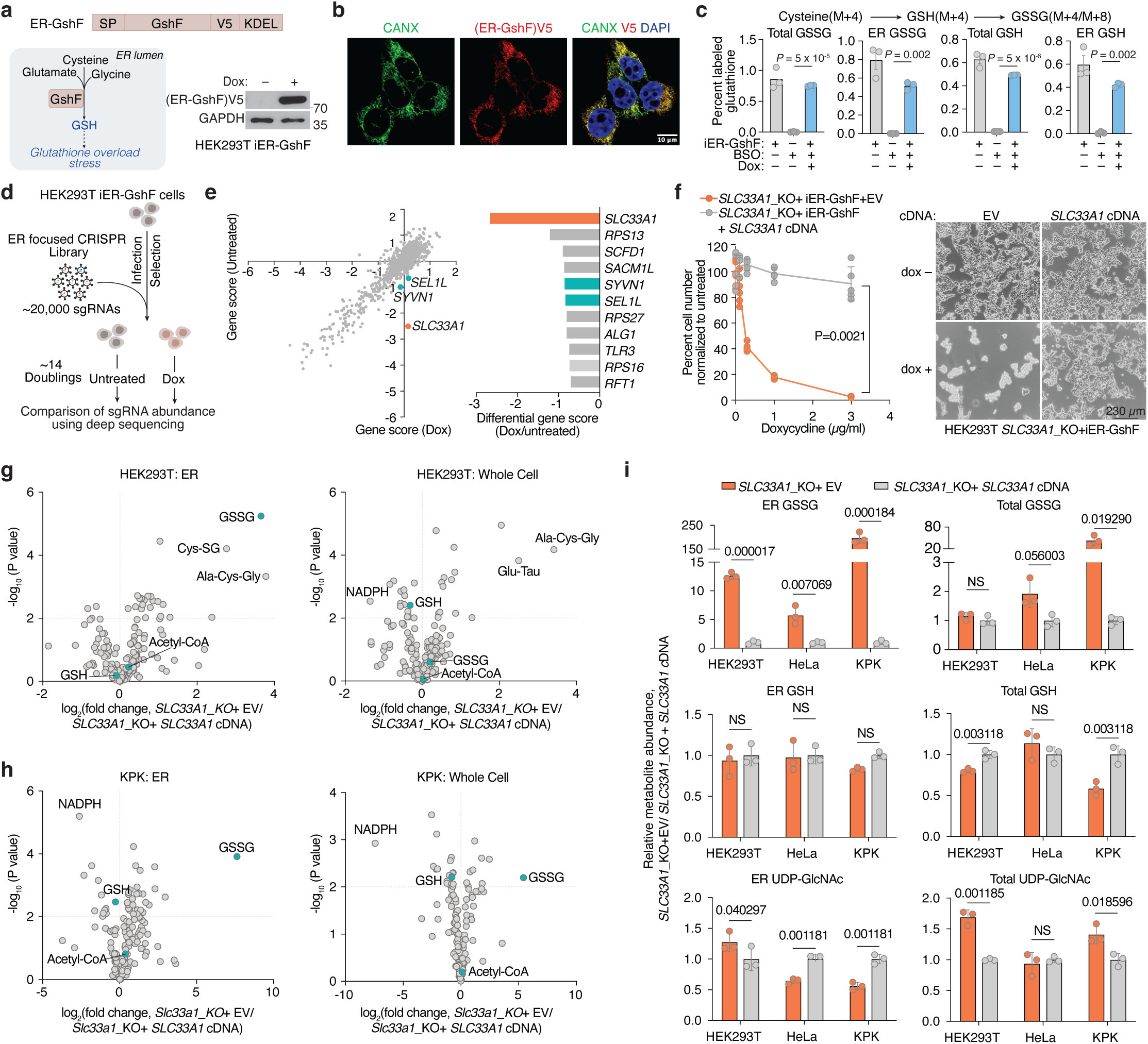
Identification of SLC33A1 as a regulator of ER glutathione homeostasis. a. Top, schematic of engineered *GshF* construct targeted to ER (*ER-GshF*), with signal peptide (SP) from ERP44 at N-terminus, V5 tag and ER retention signal KDEL at C-terminus. Bottom left, schematic of GSH synthesis in ER catalyzed by ER-GshF. Bottom right, immunoblot analysis of ER-GshF expression from HEK293T cells expressing inducible ER-GshF (iER-GshF HEK293T) treated with 1 µg/ml doxycycline for 24 hours. Immunoblots are representative of three independent experiments with similar results. Unprocessed blots are available in Source Data. b. Immunofluorescence analysis of ER-GshF (V5, red) and calnexin (CANX, green) in iER-GshF HEK293T cells treated with 1 µg/ml doxycycline for 48 hours. Micrographs are representative of two independent experiments. Immunofluorescence images are representative of three independent experiments with similar results. Source images are available in Source Data. c. Percent labeled glutathione from HEK293T cells expressing a vector control or inducible ER-GshF pre-treated with or without BSO and doxycycline. Cells pretreated with or without 1 mM BSO and 1 µg/ml doxycycline for 24 hours, were switched to cystine free media with 200 µM isotope labeled cystine (³C₆, ¹⁵N₂) for 8 hours before harvesting the cells, BSO and doxycycline were kept the same as pretreatment during labeling. Data are presented mean ± SEM. Statistical significance was assessed using a two-tailed paired Student’s t-test. n = 3 biological replicates. Source numerical data are available in Source Data. d. Schematic of the ER-focused CRISPR genetic screens in iER-GshF HEK293T cells cultured in the presence or absence of 1 µg/ml doxycycline for 14 doublings. e. Left, CRISPR gene scores in iER-GshF HEK293T cells cultured in the presence or absence of 1 µg/ml doxycycline for 14 doublings. Top scoring hits color-coded. Pearson correlation coefficient, two-sided. Right, differential gene score from iER-GshF HEK293T cells cultured with doxycycline compared to untreated. Top genes sensitizing iER-GshF HEK293T cells under doxycycline treatment are shown. Source numerical data are available in supplementary table 7. f. Left, percent cell number from *SLC33A1* knockout HEK293T cells expressing inducible ER-GshF complemented with a vector control or *SLC33A1* cDNA under different concentrations of doxycycline for 4 days. Numbers under doxycycline treated are normalized to untreated. Right, representative images of the indicated cells. Data are presented as mean ± SD. Statistical significance was assessed using a two-tailed unpaired t-test with Welch’s correction. n = 5 biological replicates. Source numerical data are available in Source Data. g. Left, volcano plot showing relative fold change (log_2_) in ER metabolite abundance versus −log (P values) from *SLC33A1* knockout ER-tag HEK293T cells expressing a vector control or *SLC33A1* cDNA. Statistical significance was determined by multiple two-tailed unpaired t-tests. The dotted line represents *P*=0.01. Right, volcano plot showing relative fold change (log_2_) in whole-cell metabolite abundance versus −log (P values) from *SLC33A1* knockout ER-tag HEK293T cells expressing a vector control or *SLC33A1* cDNA. Statistical significance was determined by multiple two-tailed unpaired t-tests. The dotted line represents *P*=0.01. n = 3 biological replicates. Source numerical data are available in Source Data. h. Left, volcano plot showing relative fold change (log_2_) in ER metabolite abundance versus −log (P values) from *Slc33a1* knockout ER-tag KPK cells expressing a vector control or *SLC33A1* cDNA. Statistical significance was determined by multiple two-tailed unpaired t-tests. The dotted line represents *P*=0.01. Right, volcano plot showing relative fold change (log_2_) in whole-cell metabolite abundance versus −log (P values) from *Slc33a1* knockout ER-tag KPK cells expressing a vector control or *SLC33A1* cDNA. Statistical significance was determined by multiple two-tailed unpaired t-tests. The dotted line represents *P*=0.01. n = 3 biological replicates. Source numerical data are available in Source Data. i. Relative metabolite abundance of indicated whole cell or ER metabolites from *SLC33A1* knockout ER-tag HEK293T, HeLa and KPK cells expressing a vector control compared to those expressing *SLC33A1* cDNA. ER UDP-GlcNAc abundance is shown to indicate ER amount. Data are presented as mean ± SD. Statistical significance was assessed using two-tailed unpaired t-tests with false discovery rate (FDR) correction using the two-stage step-up method of Benjamini, Krieger and Yekutieli (Q = 1%). n = 3 biological replicates. Source numerical data are available in Source Data.

Since SLC33A1 is predicted to function as an ER-localized small molecule transporter and is essential under conditions of GSH overload, we hypothesized that it regulates ER glutathione homeostasis via its transport activity. To test this, we utilized ER-IP coupled with LC-MS-based metabolomics to profile ER metabolites in *SLC33A1* knockout cells and in cells complemented with *SLC33A1* cDNA (Extended Data Fig. 3d). These experiments were performed in HEK293T, HeLa and murine lung cancer cell lines Kras^G12D^, p53^-/-^(KP) and Kras^G12D^, p53^-/-,^ Keap1^-/-^(KPK), the latter exhibiting elevated GSH levels due to the activation of NRF2, a master regulator of the antioxidant pathway^26^. Among all detected metabolites, GSSG showed the most pronounced accumulation in SLC33A1-deficient cells, exhibiting a specific increase in the ER (5-12-fold in HEK293T, HeLa and KP cells, 100-fold in KPK cells). Loss of *SLC33A1* did not affect GSH in the ER, whole-cell glutathione pools, or most other ER metabolites (Fig. 2g-i; Extended Data Fig. 3e-g). Of note, cellular NADPH levels were significantly reduced upon *SLC33A1* loss, potentially due to increased activity of the cytosolic thioredoxin system, which may compensate by supplying reducing equivalents for the ER at the expense of cytosolic NADPH^27,28^. Although SLC33A1 has been proposed to import acetyl-CoA into the ER lumen^25^, we did not observe any change in ER acetyl-CoA levels upon *SLC33A1* loss (Extended Data Fig. 3h). Additionally, while we observed changes in some cysteine-containing tripeptides, levels of most di– and tripeptides remained unaltered (Extended Data Fig. 3i), supporting the selectivity of SLC33A1 for GSSG. Finally, a co-evolution analysis across eukaryotes identified a strong functional association between *SLC33A1* and glutamate-cysteine ligase catalytic subunit, *GCLC*, the major glutathione synthesis enzyme (Pearson Correlation = 0.791) (Extended Data Fig. 3j), further supporting its involvement in glutathione homeostasis. These findings establish SLC33A1 as a critical regulator of ER GSSG levels and establish its central role in glutathione homeostasis in the ER.

### SLC33A1 is an ER GSSG transporter

Given the accumulation of ER GSSG upon *SLC33A1* loss, we reasoned that SLC33A1 may mediate GSSG transport across the ER membrane. To evaluate this hypothesis, we first depleted ER glutathione using BSO (Extended Data Fig. 4a) and measured the uptake of radiolabeled GSSG into ER isolated from *SLC33A1* knockout cells or those complemented with *SLC33A1* cDNA (Fig. 3a). GSSG accumulated in a time-dependent manner only in ER from *SLC33A1* expressing cells (Fig. 3a), with an apparent Michaelis constant (K_m_) of 2.4 mM (Fig. 3b). These results are consistent with the millimolar concentrations of GSSG found in the ER lumen^9,11^. We next asked whether GSSG directly binds to SLC33A1 by determining the effects of putative substrates on the thermal stability of purified SLC33A1 (Fig. 3c; Extended Data Fig. 4b). The addition of GSSG, but not GSH or glycine, resulted in marked shifts in SLC33A1 melting temperature, as assessed by measuring tryptophan fluorescence at 330 nm and 350 nm during a temperature ramp (Fig. 3b; Extended Data Fig. 4c-e). Cys-SG, which accumulates in the ER upon *SLC33A1* loss, also altered the melting temperature of SLC33A1 (Fig. 3b; Extended Data Fig. 4f). Conversely, several related metabolites whose abundances remain unchanged in cells lacking SLC33A1 had little to no effect on the melting temperature of SLC33A1, supporting selective recognition of GSSG (Extended Data Fig. 4g). In line with this, GSH and acetyl-CoA uptake into ER was unaffected by *SLC33A1* loss (Extended Data Fig. 4h, i). To directly test transport activity, we reconstituted purified SLC33A1 into lipid vesicles pre-loaded with unlabeled GSSG and measured the counterflow of radiolabeled GSSG into SLC33A1-containing and protein-free liposomes (Extended Data Fig. 4j). We observed a time-dependent accumulation of radiolabeled GSSG in the SLC33A1-containing liposomes compared to that in the empty liposomes (Fig. 3d). Moreover, unlabeled GSSG but not acetyl-CoA competitively inhibited radiolabeled GSSG transport in both ER uptake and liposome counterflow assays (Extended Data Fig. 4k-p). Collectively, these results demonstrate that SLC33A1 serves as an ER-localized GSSG transporter.

**Fig. 3:**
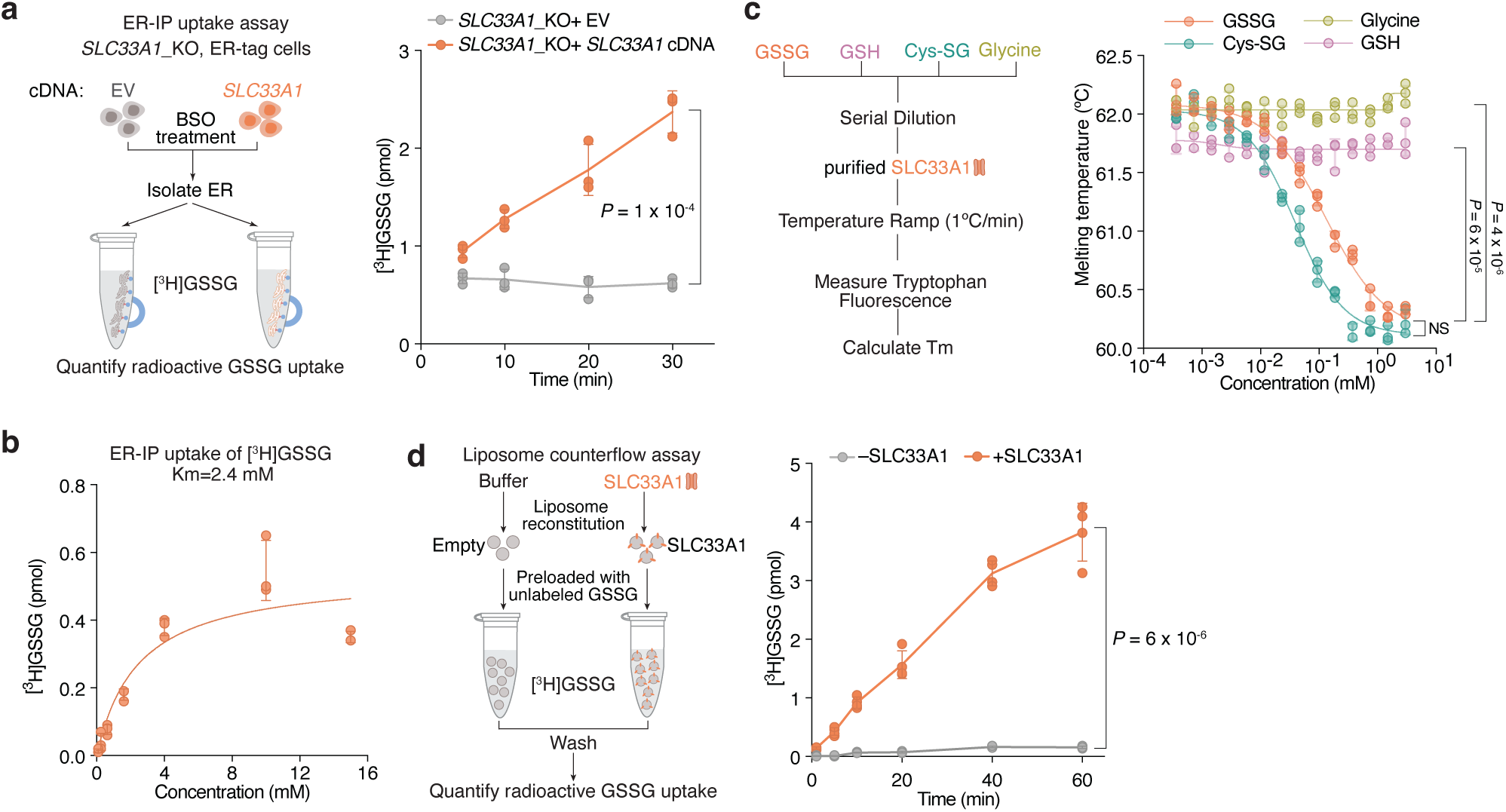
SLC33A1 is an ER GSSG transporter. a. Left, schematic depicting the experimental workflow for measuring ER GSSG uptake. ER was isolated from *SLC33A1* knockout HEK293T cells expressing vector control or *SLC33A1* cDNA after 1 mM BSO treatment for 48 hours. Right, uptake of [^3^H]GSSG into ER after incubation with 400 µM unlabeled GSSG and 4 µM radioactive GSSG at indicated times. ER was isolated from *SLC33A1* knockout HEK293T cells expressing a vector control or *SLC33A1* cDNA at indicated times. Data are presented as mean ± SD. Statistical significance was assessed using a two-tailed unpaired t-test. *n* = 3 biologically independent samples. Source numerical data are available in Source Data. b. Kinetics of [^3^H]GSSG uptake into ER was measured after 40-minute incubation with a fixed 1:500 ratio of labeled to unlabeled GSSG at the concentration of unlabeled GSSG: 0.1, 0.25, 0.64, 1.6, 4, 10, 15 mM. ER was isolated from *SLC33A1* knockout HEK293T cells expressing vector control or *SLC33A1* cDNA after 1 mM BSO treatment for 48 hours. Values represent uptake in *SLC33A1* knockout ER expressing *SLC33A1* cDNA, after subtraction of background signal from *SLC33A1* knockout ER expressing vector control. The curve was fitted to the Michaelis–Menten model using total GSSG concentration. Data are presented as mean ± SD. *n* = 3 biologically independent samples. Source numerical data are available in Source Data. c. Left, schematic depicting the workflow for thermal shift assay. GSSG, GSH, Cys-SG and glycine were serially diluted, and then incubated with purified SLC33A1. Samples were subjected to temperature ramping (1°C/min) while monitoring tryptophan fluorescence to determine melting temperatures (Tm). Right, calculated melting temperatures of SLC33A1 in the presence of GSSG (orange), Cys-SG (teal), glycine (green) and GSH (purple). Data are presented as mean ± SD. *n* = 4 technically independent samples. Source numerical data are available in Source Data. d. Left, schematic depicting of liposome-based GSSG counterflow assay. Liposomes were reconstituted with either buffer (empty) or purified SLC33A1, preloaded with unlabeled GSSG, followed by incubation with [^3^H]GSSG. After washing, radioactive GSSG uptake was quantified. Right, counterflow transport of [^3^H]GSSG into empty or SLC33A1 reconstituted liposomes preloaded with 5 mM unlabeled GSSG, followed by incubation with 5 µM [^3^H]GSSG for indicated times. Data are presented as mean ± SD. Statistical significance was assessed using two-tailed unpaired t-tests with false discovery rate (FDR) correction using the two-stage step-up method of Benjamini, Krieger and Yekutieli (Q = 1%). *n* = 4 technically independent samples. Source numerical data are available in Source Data.

### Mechanism of GSSG transport by SLC33A1

To determine how GSSG is recognized and transported by SLC33A1, we collected cryogenic electron microscopic (cryo-EM) images of human SLC33A1 vitrified in the presence of 20 mM GSSG. However, due to its small size, we were unable to calculate a reconstruction from the particle images of SLC33A1 alone that was suitable for model building. Fiducial markers such as antibody fragments that bind to ordered epitopes are routinely used to increase the size of small membrane protein targets and provide extramembranous features to aid image alignment^29–31^. We therefore introduced the MAP-tag, an 8-residue tag with the sequence ‘GDGMVPPG’, into 3 locations in SLC33A1 to enable binding of a synthetic antibody fragment^32^. Among these, we successfully expressed and purified the construct in which the MAP was inserted into the loop connecting TM9 and TM10 (SLC33A1^MAP^) (Extended Data Fig. 5a-c). Expression of SLC33A1^MAP^ was sufficient to rescue the anti-proliferative effect of glutathione overload in SLC33A1 knockout cells, confirming that SLC33A1^MAP^ is functional (Extended Data Fig. 5d). Cryo-EM analysis of SLC33A1^MAP^ in complex with Pmab-1 Fv-clasp that specifically recognizes the MAP-tag^32,33^ revealed that the SLC33A1^MAP^-Fv-clasp complex can adopt several conformations in the presence of GSSG (Extended Data Fig. 6a). Iterative classification of the best resolved 3D class identified a subset of particles that yielded a 3.2 Å reconstruction of the SLC33A1^MAP^-Fv-clasp complex (Fig. 4a; Extended Data Fig. 6a-d and Supplementary Table 4). Focused refinement using a mask encompassing the transmembrane domain of SLC33A1^MAP^ improved the interpretability of the reconstruction and enabled us to build and refine a model of SLC33A1 containing residues 70-282 and 293-550, which correspond to residues 70-282 and 293-547 in wild-type SLC33A1 (Fig. 4b, c; Extended Data Fig. 6e-j and Supplementary Table 4). For simplicity, we use the wild-type residue numbering when discussing the structure of SLC33A1^MAP^. SLC33A1^MAP^ is comprised of two 6-helix domains that are connected by a poorly ordered linker that is not modelled in the structure (Fig. 4b, c). Similar to other members of the Major Facilitator Superfamily, the two repeats of SLC33A1^MAP^ share considerable structural homology as evidenced by a main-chain RMSD of 4.6 Å between TM1-TM6 and TM7-TM12^34^. The two repeats are arranged around a central pseudo-two-fold axis and create a central cavity continuous with the cytosol that extends more than 40 Å into the transporter. The luminal end of the cavity is sealed off by the luminal ends of TM1, TM2, TM7 and TM8 (Fig. 4c, d).

**Fig. 4:**
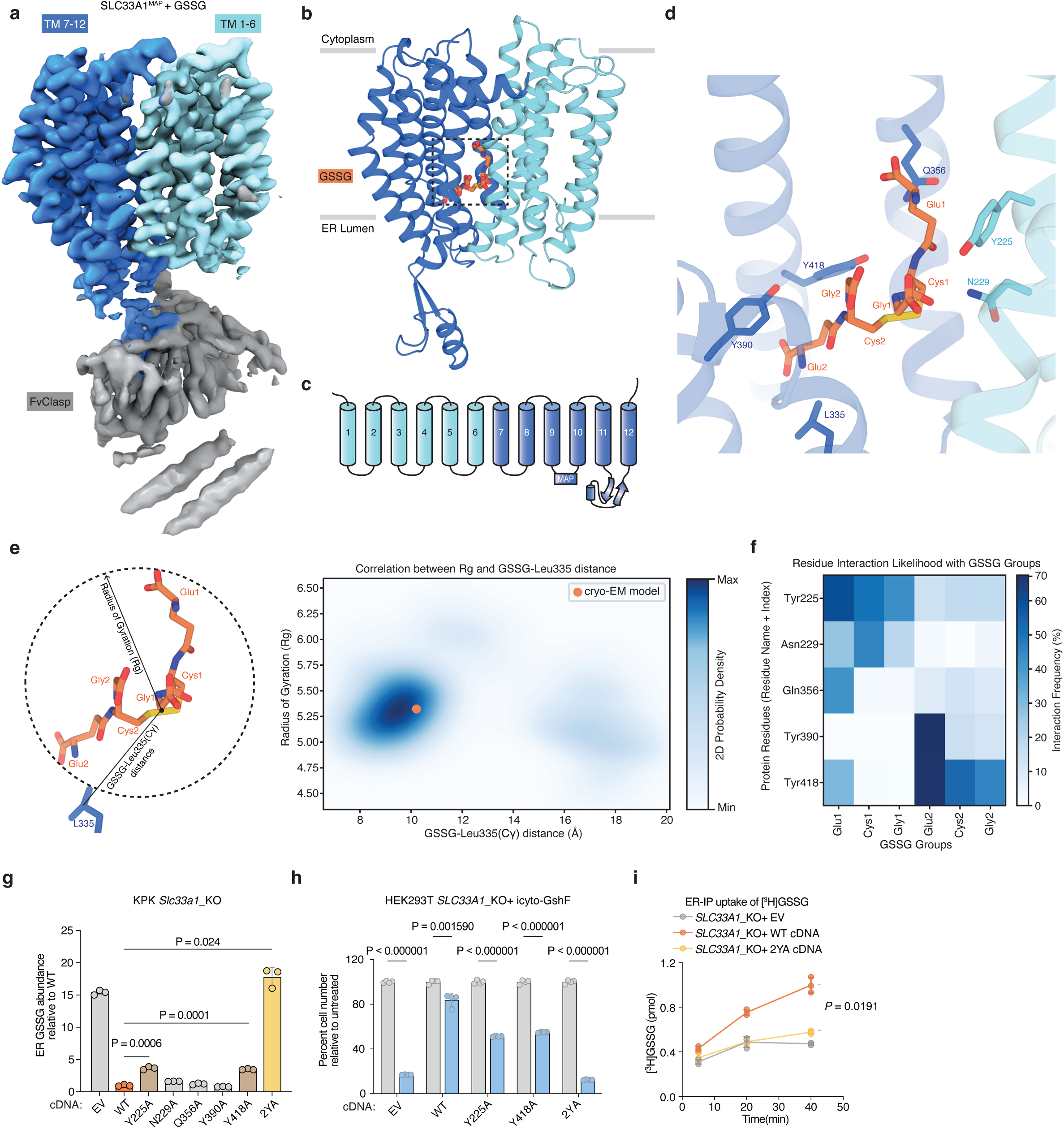
Structure of SLC33A1. a. Consensus reconstruction of SLC33A1^MAP^ in the presence of PMab-1 Fv-clasp and GSSG. b. Atomic model of GSSG-bound SLC33A1^MAP^ in a cytosolic-facing conformation derived from TMD focus refinement reconstruction. Transmembrane helices 1-6 and 7-12 are shown in the light blue and dark blue, respectively. GSSG is shown in orange stick representation. c. Schematic of SLC33A1 topology colored as in panel b. d. GSSG-binding site of SLC33A1. Side chains that comprise the substrate-binding site are shown as sticks. e. Left, schematic of GSSG with depictions of measurements of the radius of gyration (Rg) and distance between GSSG center of mass and Leu335 Cγ. Right, 2D probability density plot showing the correlation between GSSG (Rg) and distance between GSSG center of mass and Leu335 Cγ during MD simulations. Cryo-EM pose is shown as an orange circle. Source numerical data are available in Source Data. f. Heat map showing the interaction frequency between residues of GSSG and the residues of SLC33A1 that comprise the substrate binding pocket during MD simulations. Source numerical data are available in Source Data. g. Relative GSSG abundance in ER isolated from *Slc33a1* knockout KPK cells expressing wild-type, mutant *SLC33A1* cDNAs or a vector control. Data are presented as mean ± SD. Statistical significance was assessed using a two-tailed paired t-test. n = 3 biological replicates. Source numerical data are available in Source Data. h. Percent cell number from *SLC33A1* knockout HEK293T cells expressing inducible cyto-GshF (icyto-GshF) complemented with a vector control, WT or mutant *SLC33A1* cDNAs in the presence or absence of 1 µg/ml doxycycline treatment for 4 days. Numbers under doxycycline treated are normalized to untreated. Data are presented as mean ± SD. Statistical significance was assessed using two-tailed unpaired t-tests with false discovery rate (FDR) correction using the two-stage step-up method of Benjamini, Krieger and Yekutieli. *n* = 4 biologically independent samples. Source numerical data are available in Source Data. i. Uptake of [^3^H]GSSG into ER after incubation with 80 µM unlabeled GSSG and 10 µM [^3^H]GSSG at indicated times. ER was isolated from *Slc33a1* knockout KPK cells expressing WT, 2YA mutant *SLC33A1* cDNAs or a vector control at indicated times. Cells were treated with 1 mM BSO for 48 hours before ER isolation. Data are presented as mean ± SD. Statistical significance was assessed using a two-tailed paired t-test. *n* = 3 biologically independent samples. Source numerical data are available in Source Data.

Within the central cavity, we observed a non-protein density into which we modelled one GSSG molecule (Fig. 4d; Extended Data Fig. 6k). The bound GSSG adopts an extended conformation that reaches from the narrow luminal end of the cavity where one of the glutamate residues (Glu2) is surrounded by Leu 335, Tyr390 and Tyr418 to the middle of the cavity where the opposing glutamate residue (Glu1) contacts Tyr225, Gln356 and Tyr426. Despite GSSG having a net charge of –2, we do not observe GSSG forming strong ionic interactions with any cavity lining residues. Instead, the cavity is lined with an abundance of aromatic residues that contact GSSG including Tyr225, Tyr390, Tyr418, Tyr426, and Tyr429. These aromatic residues shape the cavity to allow GSSG to bind and are positioned to form specific interactions with GSSG. For example, Tyr225 is positioned such that it may form a cation-π interaction with the backbone nitrogen atom of Glu1. We next performed a series of all-atom molecular dynamics simulations initiated from the cryo-EM model and investigated the interactions between GSSG and SLC33A1^MAP^ (Extended Data Fig. 7a and Supplementary Table 5 and 6). To assess the conformation of GSSG during the simulations, we measured its radius of gyration (measurement of the spatial distribution of the molecule around its center of mass), which is highly dynamic in solution^35,36^, and its distance from the Cγ of the relatively static Leu335 (Fig. 4e; Extended Data Fig. 7b, c). Across 3.4 µs of aggregated simulation time, we observed that GSSG adopted a primary pose that closely resembles the extended conformation present in the cryo-EM structure. GSSG primarily interacts with aromatic residues during the simulations, with Tyr225, Tyr390, and Tyr418 being among the most contacted residues. In addition to the aromatic residues, GSSG also commonly interacts with Asn229 and Gln356 (Fig. 4f; Extended Data Fig. 7d).

To assess the functional role of these substrate-binding residues, we measured GSSG abundance in ER from *Slc33a1* knockout cells expressing wild-type, mutant *SLC33A1* cDNAs or an empty vector. While individual mutation of Y225 or Y418 to alanine led to a moderate increase in GSSG levels compared to wild type SLC33A1, the double mutant (2YA) phenocopied GSSG accumulation seen in *Slc33a1* knockout cells, indicating that Y225 and Y418 cooperate to coordinate GSSG (Fig. 4g; Extended Data Fig. 8a). Consistent with these results, a comparison of the structures of SLC33A1 in the presence of GSSG or acetyl-CoA reveals that the substrates may bind to overlapping residues including Y225 and Y418^37^ (Extended Data Fig. 8b, c). In line with these results, expression of wild type or single mutant cDNAs fully or only partially rescued cell death of *SLC33A1* knockout cells in response to glutathione overload, whereas the 2YA mutant failed to confer protection (Fig. 4h). Additionally, 2YA mutant completely blocked GSSG uptake into ER (Fig. 4i) and reduced its transport by 50% in liposome counterflow assays compared to wild type SLC33A1 (Extended Data Fig. 8d). Approximately 5-fold more GSSG was needed to shift the midpoint of the melting curve of the Y2A mutant compared to wild-type SLC33A1, suggesting that its diminished transport activity was due to its reduced affinity for GSSG (Extended Data Fig. 8e). Collectively, these observations support a model where GSSG adopts an extended conformation when bound to SLC33A1 that is stabilized by numerous interactions with cavity-lining aromatic residues.

Finally, variants in SLC33A1 have been linked to Huppke-Brendel syndrome, a rare autosomal recessive neurodegenerative disorder^38^. To evaluate the functional impact of these variants, we expressed four of the patient-derived variants with single amino substitutions in *Slc33a1* knockout cells and measured ER GSSG levels. Among the disease variants investigated, the A110P variant was non-functional, phenocopying the GSSG accumulation seen in *Slc33a1* knockout cells (Extended Data Fig. 8f). Mapping the disease-associated mutations onto the GSSG-bound structure revealed that A110 is located at the TM2–TM11 interface (Extended Data Fig. 8g, h). In MFS transporters, the conformation of the TM2-TM11 interface rearranges during the transport cycle to enable alternating access to the substrate-binding site^34^, suggesting that the A110P variant may be due to diminished transport efficacy.

### Cellular outcome of ER-specific redox dysfunction in human cells

We next sought to determine whether *SLC33A1* loss and the resulting GSH/GSSG imbalance would perturb cysteine reactivity of proteins folded in ER and adversely impact ER function. Indeed, a systematic profiling of oxidized cysteine reactivity in *Slc33a1* knockout cells compared to cDNA-complemented cells revealed increased cysteine oxidation in proteins of secretory pathway compartments, but not of other organelles (Fig. 5a, b; Extended Data Fig. 9a). Notably, several protein disulfide isomerases (PDIs) exhibited increased cysteine oxidation, including non-catalytic (Trx-like) domains of PDIA3 (C^85^X_6_C^92^), and C270 of DNAJC10 (C^265^X_5_C^270^), as well as the regulatory cysteine of the ER oxidase ERO1A (C^130^) (Fig. 5c; Extended Data Fig. 9b). PDIs facilitate disulfide oxidation, isomerization, and reduction, processes tightly regulated by ER redox balance. PDIs catalyzing disulfide oxidation are primarily maintained in oxidized states by ERO1^39^, whereas those involved in reduction or isomerization, processes essential for resolving non-native disulfides during oxidative folding and for reducing disulfides prior to protein degradation, are typically kept in reduced states, likely through direct interaction with GSH^40^. For example, PDIA3 and DNAJC10, which mediate disulfide isomerization and reduction, respectively, require reduced states to function effectively^41–43^. Indeed, AMS modification immunoblots confirmed that PDIs and ERO1A, which normally have reduced forms, are more oxidized in *Slc33a1* knockout cells than controls, whereas PDIA1, which is normally oxidized, remains unchanged upon *Slc33a1* loss (Fig. 5d; Extended Data Fig. 9c), an effect rescued by BSO treatment (Extended Data Fig. 9d) but not the transport-defective 2YA mutant (Extended Data Fig. 9e). Consistent with *in vitro* evidence that glutathione directly reduces PDIA3 at physiological concentrations^44^, our findings suggest that glutathione contributes net reducing equivalents to the ER to maintain some PDIs in a reduced state, a process disrupted by GSSG accumulation

**Fig. 5:**
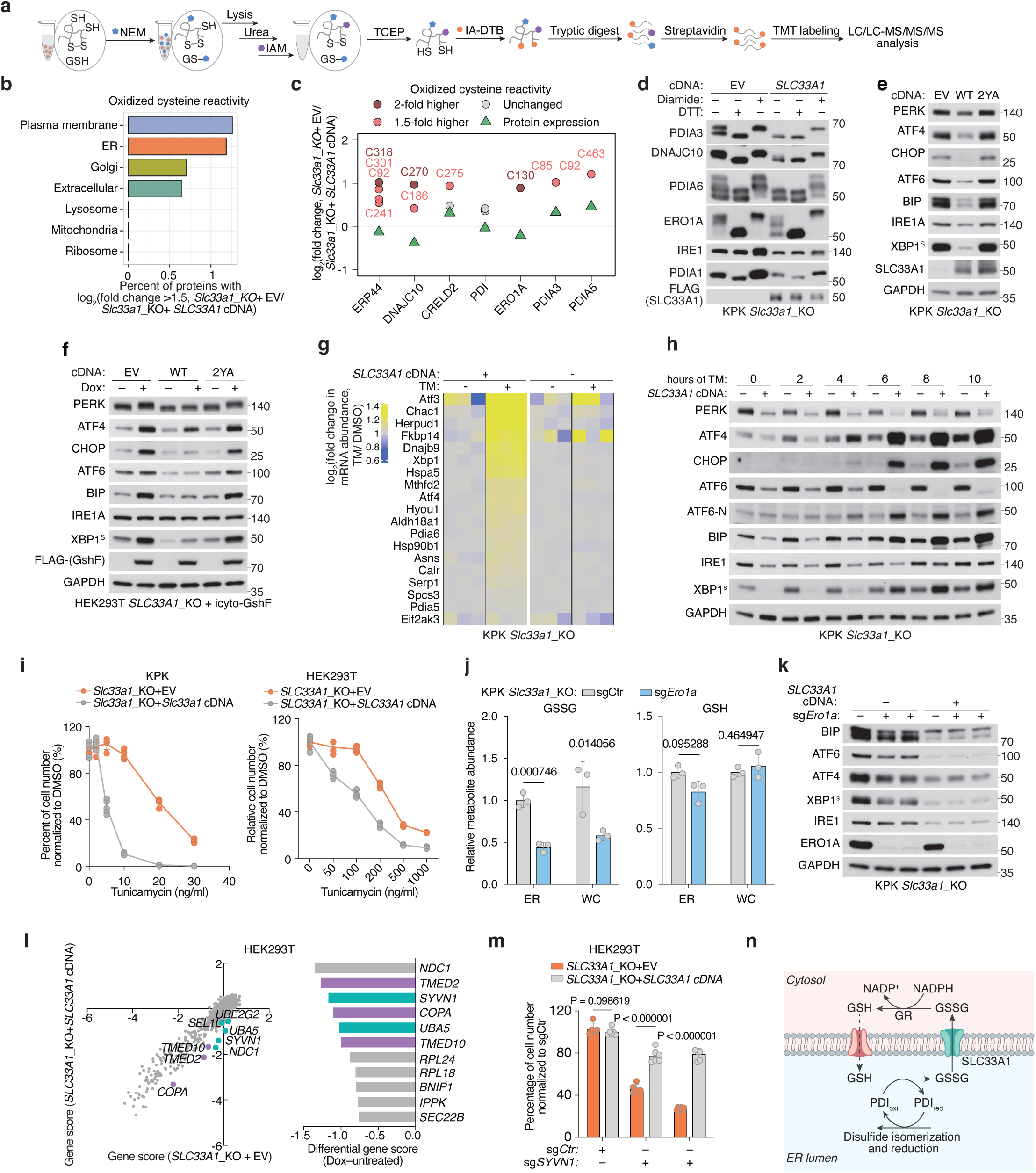
Cellular outcome of ER-specific redox dysfunction in mammalian cells. a. Schematic depicting the chemical proteomic workflow for profiling oxidized cysteines. Cells are treated with N-ethylmaleimide (NEM) before lysis and Iodoacetamide (IAM) after lysis to alkylate the reduced cysteines and GSH. Oxidized cysteines are reduced by Tris(2-carboxyethyl) phosphine (TCEP) before being labeled with iodoacetamide-desthiobiotin (IA-DTB) and enrichment via streptavidin agarose beads for further tandem mass tag (TMT) labeling and LC/LC-MS/MS/MS analysis. b. Bar graph representing the percent of proteins within indicated organelles showing higher than 1.5-fold difference (log_2_) in reactivity of oxidized cysteines from *Slc33a1* knockout KPK cells expressing a vector control compared to those expressing *SLC33A1* cDNA. Source numerical data are available in Source Data. c. Fold change (log_2_) in oxidized cysteine reactivity of indicated proteins from *Slc33a1* knockout KPK cells expressing a vector control compared to those expressing *SLC33A1* cDNA. Dark red and light red mark cysteine residues with ≥2-fold and ≥1.5-fold reactivity changes compared to protein expression, respectively. Grey marks unchanged cysteines. Green marks protein expression. Data are median values for n = 2 (or more) independent cysteine reactivity profiling experiments and n = 1 independent unenriched proteomics experiments. Each biological independent experiment had 2-3 technical replicates for each condition. Source numerical data are available in Source Data. d. Immunoblot analysis of 4-acetamido-4′-maleimidylstilbene-2,2′-disulfonic acid (AMS) modified proteins from *Slc33a1* knockout KPK cells expressing a vector control or *SLC33A1* cDNA. Cells were reduced with 10 mM DTT or oxidized with 5 mM diamide for 5 minutes prior to treatment with 25 mM NEM for 10 minutes before lysis. The bands of oxidized and reduced proteins are labeled. Immunoblots are representative of four independent experiments with similar results. Unprocessed blots are available in Source Data. e. Immunoblot analysis of indicated proteins from *Slc33a1* knockout KPK cells expressing a vector control (EV), WT or 2YA mutant *SLC33A1* cDNAs. Immunoblots are representative of three independent experiments with similar results. Unprocessed blots are available in Source Data. f. Immunoblot analysis of indicated proteins from *SLC33A1* knockout HEK293T cells expressing an inducible cyto-GshF (icyto-GshF) complemented with a vector control WT or 2YA mutant *SLC33A1* cDNA treated with 1 µg/ml doxycycline for 48 hours. Immunoblots are representative of three independent experiments with similar results. Unprocessed blots are available in Source Data. g. Heatmap of fold change (log_2_) in unfolded protein response (UPR) target mRNAs from *Slc33a1* knockout KPK cells expressing a vector control or *SLC33A1* cDNA comparing cells treated with tunicamycin to DMSO. Cells were treated with DMSO or 10 ng/ml tunicamycin (TM) for 8 hours before RNA extraction. Source numerical data are available in Source Data. h. Immunoblot analysis of indicated proteins from *Slc33a1* knockout KPK cells expressing a vector control or *SLC33A1* cDNA. Cells were treated with 20 ng/ml tunicamycin for indicated time. Immunoblots are representative of four independent experiments with similar results. Unprocessed blots are available in Source Data. i. Left, percent of cell number from *Slc33a1* knockout KPK cells expressing a vector control or *SLC33A1* cDNA treated with DMSO or tunicamycin for 4 days under indicated concentrations. n = 5 biological replicates. P < 0.0001. Right, percent of cell number from *SLC33A1* knockout HEK293T cells expressing a vector control or *SLC33A1* cDNA treated with DMSO or tunicamycin for 4 days under indicated concentrations. n = 4 biological replicates. P< 0.0001. Numbers under tunicamycin treatment were normalized to DMSO untreated. Data are presented as mean ± SD. Statistical significance was assessed using two-tailed unpaired t-tests with false discovery rate (FDR) correction using the two-stage step-up method of Benjamini, Krieger and Yekutieli (Q = 1%). Source numerical data are available in Source Data. j. Relative metabolite abundance of indicated whole cell or ER metabolites from *Slc33a1* knockout KPK cells expressing a sgRNA targeting intergenic region (sgControl) or two different sgRNAs targeting *Ero1la*. Data are presented as mean ± SD. Statistical significance was assessed using two-tailed unpaired t-tests with false discovery rate (FDR) correction using the two-stage step-up method of Benjamini, Krieger and Yekutieli (Q = 1%). n = 3 biological replicates. Source numerical data are available in Source Data. k. Immunoblot analysis of indicated proteins from *Slc33a1* knockout KPK cells complemented with a vector control or *SLC33A1* cDNA expressing a sgRNA targeting intergenic region (sgControl) or two different sgRNAs targeting Ero1la. Immunoblots are representative of three independent experiments with similar results. Unprocessed blots are available in Source Data. l. Left, CRISPR gene scores in *SLC33A1* knockout HEK293T cells expressing a vector control or *SLC33A1* cDNA cultured for 14 doublings. Top scoring hits color-coded. Pearson correlation coefficient, two-sided. Right, top-scoring genes that sensitize cells to *SLC33A1* loss. Source numerical data are available in supplementary table 8. m. Percent of cell number from *SLC33A1* knockout HEK293T cells complemented with a vector control or *SLC33A1* cDNA expressing a sgRNA targeting intergenic region (sgControl) or two different sgRNAs targeting *SYVN1* after 4-day proliferation. Numbers of sg*SYVN1* were normalized to sgCtr. Data are presented as mean ± SD. Statistical significance was assessed using two-tailed unpaired t-tests with false discovery rate (FDR) correction using the two-stage step-up method of Benjamini, Krieger and Yekutieli (Q = 1%). n = 6 biological replicates. Source numerical data are available in Source Data. n. Schematic of GSSG export by SLC33A1 to maintain ER redox homeostasis. GSH imported into ER is oxidized to GSSG upon reducing PDI family proteins that catalyze disulfide reduction and isomerization. SLC33A1 exports GSSG into cytosol, where it is reduced back to GSH by glutathione reductase (GR).

Given the essential role of PDIs in facilitating correct disulfide bond formation, we hypothesized that their oxidation disrupts this process and activates the unfolded protein response (UPR). To test this, we performed RNA sequencing of *Slc33a1* knockout and cDNA-complemented cells treated with tunicamycin, a classical UPR inducer that blocks protein N-glycosylation. Consistent with previous studies^45,46^, *Slc33a1* loss activated the UPR (Extended Data Fig. 10a, b), indicating that GSSG accumulation leads to accumulation of misoxidized and misfolded proteins. While wild-type *SLC33A1* was sufficient to alleviate UPR in *SLC33A1* knockout cells under normal conditions and during GSH overload, the transport-defective 2YA mutant failed to do so (Fig. 5e, f), confirming that GSSG export activity is required for SLC33A1 to maintain ER proteostasis. Notably, disease-associated variants A110P, Y366A and Y377A also exhibited elevated UPR activation (Extended Data Fig. 10c). Unexpectedly, *SLC33A1* loss attenuated tunicamycin-induced UPR activation in KPK cells (Fig. 5g, h; Extended Data Fig. 10d-f). This paradoxical effect of PDI oxidation highlights a dual role: PDI oxidation disrupts disulfide bond reduction and isomerization, leading to protein misfolding and chronic UPR activation. However, in response to acute UPR induction, PDI oxidation attenuates further UPR activation, potentially preserving cell viability by mitigating excessive stress responses. Consistent with this, *SLC33A1* loss enhanced resistance to tunicamycin-induced cell death (Fig. 5i; Extended Data Fig. 10g). This observation also aligns with prior evidence showing that PDIs regulate UPR via redox modulation of the luminal domains of the UPR receptors, such as dimerization^47–49^. Given that the PDIA1-ERO1 pathway is the primary driver of disulfide oxidation^39^ and GSSG production is downstream of disulfide oxidation (Extended Data Fig. 10h), we next asked whether modulating ERO1 activity could influence GSSG accumulation and UPR activation. *ERO1A* loss decreased ER GSSG levels and attenuated UPR activation in *Slc33a1* knockout cells (Fig. 5j, k). Remarkably, despite this redox stress, *SLC33A1*-deficient cells maintained their proliferation, suggesting compensatory pathways mitigate ER-specific oxidative stress. To identify such pathways, we performed CRISPR-Cas9 genetic screens in *SLC33A1* knockout cells and cDNA complemented cells using ER-focused sgRNA library. Top hits included genes involved in COPI-mediated retrograde transport from the Golgi to the ER (*TMED2*, *TMED10*, *COPA*), which likely retrieve misfolded proteins for degradation, as well as key components of ER protein quality control pathways, including the ER-associated degradation E3 ligase *SYVN1*, and the UFMylation pathway E1 enzyme *UBA5* (Fig. 5l). These results indicate that protein quality control mechanisms operating at multiple stages of the secretory pathway are essential for clearing misfolded proteins and maintaining cell survival. Consistently, SYVN1 loss was lethal in S*LC33A1* knockout cells but not in cDNA complemented counterparts (Fig. 5m; Extended Data Fig. 10i). Our findings demonstrate that SLC33A1-mediated GSSG export is essential for maintaining ER glutathione homeostasis, supporting PDI function, regulating protein cysteine reactivity, and enabling cellular adaptation to oxidative stress (Fig. 5n; Extended Data Fig. 10h).

## Discussion

Glutathione availability has long been proposed to maintain ER redox homeostasis and support protein maturation. One mechanism by which the ER sustains this redox balance is through glutathione transport, as the ER lacks a glutathione reductase to regenerate GSH from GSSG internally. Here, we identify SLC33A1 as the primary ER transporter for oxidized glutathione (GSSG), facilitating its export down its concentration gradient to the cytosol and preserving an optimal oxidative environment within the ER. Loss of *SLC33A1* disrupts this critical balance, leading to ER-specific oxidative stress and impaired protein folding. While *in vitro* studies proposed that glutathione promotes disulfide bond formation^50,51^, our findings support an alternative model, consistent with previous observations^43,44,52^, in which glutathione supplies reducing equivalents to the ER and the GSH/GSSG ratio maintains PDIs involved in disulfide isomerization and reduction in a functional, reduced state. Consistent with its role in GSSG export, our data, together with previous studies^45,46^, show that *SLC33A1* loss exacerbates ER stress across diverse cellular contexts. We also uncover a regulatory coupling between GSH/GSSG balance and ERO1, the primary oxidizing enzyme in the ER. Hyperoxidation of ER upon *SLC33A1* loss inactivates ERO1 by oxidizing its regulatory disulfide bonds (Figure 5c, d), which limits disulfide bond formation and, in turn, reduces downstream GSH oxidation and GSSG accumulation (Extended Data Fig. 10h). Together, these findings reveal a tightly coordinated system in which GSSG export, ER redox balance, and ERO1 activity are jointly regulated to maintain ER proteostasis.

While our data indicate that GSSG is the primary physiological substrate of SLC33A1, the transporter may also recognize additional substrates. Although SLC33A1 has been proposed to transport acetyl-CoA^25,37^, neither our metabolomic profiling nor ER acetyl-CoA uptake assays support this role in cultured cells. Nevertheless, we leave open this possibility that this may occur in specific contexts, warranting further biophysical and physiological studies. Instead, we observed changes in cysteine-containing di-and tripeptides upon *SLC33A1* loss (Extended Data Fig. 3i), though their functional relevance remains unclear. More broadly, our findings provide a framework for identifying additional ER metabolite transporters. Despite the presence of numerous metabolites within the ER (Figure 1i), their functional and physiological roles remain poorly defined. Uncovering the specific carriers for these metabolites will be essential to understanding ER biology.

Mutations in *SLC33A1* have been linked to Huppke-Brendel disease^38^, a rare neurodevelopmental disorder characterized by severe intellectual disability, motor deficits, and progressive neurodegeneration. Our data reveal that disease-associated variants of SLC33A1 impair GSSG export and activate UPR, raising the possibility that *SLC33A1* dysfunction may disrupt ER redox balance and protein folding during brain development. Indeed, ER dysfunction is a well-established contributor to neurodegenerative and developmental disorders^53^, implicating ER glutathione dysregulation may underlie broader aspects of disease pathogenesis. These insights open avenues for therapeutic intervention, including strategies to reduce ER glutathione overload through glutathione synthesis inhibitors or compounds that dissipate accumulated GSSG. Additionally, recent work highlighted an essential role for *SLC33A1* in Keap1 mutant lung cancers^45,54^, where NRF2, the master regulator of antioxidant metabolism, is constitutively activated. While NRF2 drives many downstream effector pathways for boosting antioxidant functions, ER GSSG accumulation likely drives the dependency of Keap1 mutant cancer cells on *SLC33A1*. Targeting ER redox homeostasis may thus represent a viable therapeutic approach in cancers characterized by glutathione accumulation. Collectively, our work establishes SLC33A1 as a key regulator of ER glutathione export with implications for both neurodevelopmental disease and cancer. Future studies will be needed to define its physiological roles across diverse tissues and pathological contexts.

## Acknowledgements

We thank John Chodera for assistance with the molecular dynamics simulations, M. J. de la Cruz for help with data acquisition, Olga Boudker, Melinda Diver, Krishna Reddy, and Vishnu Ganesh Ghani for advice in liposome assays, the Memorial Sloan Kettering Cancer Center HPC group for assistance with data processing and all members of the Birsoy and Hite labs for helpful suggestions. Data were generated and/or analyzed by the Proteomics Resource Center (RRID: SCR_017797), Electron Microscopy Resource Center, Bioinformatics Resource Center, Fisher Drug Discovery Resource Center, and the Flow Cytometry Resource Center at The Rockefeller University. S.L. was supported by Human Frontier Science Program (HFSP) Fellowship *LT0039/2023-L* (S.L.) (DOI: <u>10.52044/HFSP.LT00392023-L.pc.gr.169147</u>) and Robertson Therapeutic Development Fund/Kellen Women’s Entrepreneurship Fund (TDF/ WEF). V.B. is supported by the MERIT Sawyers Fellowship. Work in the Fedorova lab is supported by ‘‘Sonderzuweisung zur Unterstützung profilbestimmender Struktureinheiten’’ by the SMWK to TUD, TG70 by Sächsische Aufbaubank and SMWK, the measure is co-financed with tax funds on the basis of the budget passed by the Saxon state parliament (to M.F.), Deutsche Forschungsgemeinschaft (FE 1236/5-1, FE 1236/8-1 to M.F.), and Bundesministerium für Bildung und Forschung (031L0315A, DEEP_HCC and 01EJ2205A, FERROPath to M.F.). H.K. was supported by Pels Family Center Fellowship. E.V.V. was supported by Robertson Foundation, The Achelis and Bodman Foundation, and Searle Scholar Program. Some analyses were performed on computational resources from the Rockefeller University High Performance Computing Resource Center, RRID: SCR_025889. R.K.H was supported by the Pershing Sohn Square Prize and NIH-National Cancer Institute Cancer Center Support Grant P30-CA008748. K.B. and E.V.V. were supported by the Stavros Niarchos Foundation (SNF) as part of its grant to the SNF Institute for Global Infectious Disease Research at The Rockefeller University. K.B. was supported by NIDDK (R01 DK140337), NCI (R01 CA273233), The G. Harold and Leila Y. Mathers Charitable Foundation and is a Chan Zuckerberg Biohub Investigator.

## Author Contributions Statement

K.B. and S.L. initiated the project and K.B., S.L., M.G., R.K.H. designed the experiments. S.L. performed most of the experiments except structure related ones with the assistance of C.L. and K.W. M.G. performed structure, biochemical, and liposome related experiments under supervision of R.K.H. with the help of S.L. C.L. helped with ER immunopurification and immunoblots. K.C. and G.J.P. performed metabolomic data acquisition and analysis. Y.L. performed immunofluorescence of ER-GshF localization and helped with immunoblots. V.B. and M.G. performed molecular dynamics simulation analysis. A.M. performed proteomic analysis and all the RNA-Seq analysis. H.K., H.S. and E.V.V. performed cysteine reactivity data acquisition and analysis; M.W. and M.F. performed lipidomic data acquisition and analysis. L.U. provided structural analysis consultation. K.B., S.L., M.G., R.K.H. wrote the manuscript with input from Y.L., E.V.V., L.U., V.B., H.S., A.M.

## Methods

All experiments were performed in accordance with relevant institutional biosafety regulations and were approved by the Institutional Biosafety Committee of The Rockefeller University.

## Cell lines and reagents

Human cells lines HeLa, HEK293T cells were purchased from the ATCC. Mouse cell lines KP1 and KPK were a gift from Dr. Thales Papagiannakopoulos. Cell lines were verified to be free of mycoplasma contamination and the identities of all were authenticated by STR profiling. All the cells were maintained in DMEM media (Gibco) containing 4.5 g/L glucose, 110 mg/L pyruvate, 4mM glutamine, 10% fetal bovine serum, 1% penicillin and streptomycin unless otherwise indicated.

Antibodies against GAPDH (GTX627408) from GeneTex; GM130 (SI-9001978) from BD BioSciences, PEX19 (ab137072), SLC25A12 (ab200201) from Abcam, RPS6 (2217S), CDH1 (14472S), LAMP1(9091P), PDI (3501), CANX (2679S), CALR (12238P), citrate Synthase (14309S), PDI (3501P), IRE1 (3294), PDIA3 (2881T), PERK (5683), BiP (3177), ATF6 (65880S), CHOP (2895), ATF4(11815S), XBP-1s (40435S), SYVN1 (14773S), HA (2367S), HA (3724S), EEA1 (3288P) from Cell Signaling Technology; ATF6-N (A0202) from ABClonal; V5 (V8012-50UG), FLAG (F1804-200UG) from Sigma; DNAJC10 (13101-1-AP), PDIA6 (18233-1-AP) from Proteintech; ERO1A (sc-365526) from Santa Cruz Biotechnology; DNAJC10 (13101-1-AP), PDIA6 (18233-1-AP), TMX3 (21040-1-AP) from Proteintech. Anti-mouse IgG-HRP linked (7076S, 1:5000) and Anti-rabbit IgG-HRP linked (7074S, 1:5000) from Cell Signaling Technology. Antibodies for immunofluorescence staining were donkey anti-rabbit Alexa Fluor 488 (A21206, 1:500) and goat anti-mouse Alexa Fluor 568 (A10037, 1:500) from ThermoFisher. SLC33A1(generated in this paper).

Reagents: Radioactive GSSG (MT1003322) from Moravek Biochemicals Inc, Molar Activity by LC-MS: 14.2 Ci/mmol, concentration: 1.0 mCi/ml; 43.21 pg/ml, Packaged in: Ethanol: water (1:1) solution. ^3^C_6_, ^15^N_2_ L-cystine (CNLM-4244-H-MG-PK) from Cambridge Isotope Laboratories. Anti-HA magnetic beads (Thermo Scientific Pierce 88837), DAPI (D1306), 4-Acetamido-4’-Maleimidylstilbene-2,2’-Disulfonic Acid (AMS, A485), N-ethylmaleimide (NEM, 23030) from ThermoFisher Scientific. Polybrene (H9268), puromycin (P8833), diamide (D3648), doxycycline hyclate (D9891), iodoacetamide (IAM, I1149), tris (2 carboxyethyl) phosphine (TCEP, C4706) from Sigma. L-Buthionine-(S, R)-Sulfoximine (BSO, 14484), thapsigargin (10522) from Cayman Chemical; blasticidin from Invivogen (ant-bl-1). Iodoacetamide-desthiobiotin (IA-DTB, HY-150230) from MedChemExpress; antimycin A (ALX-380-075) from Enzo Life Sciences; tunicamycin (3516) from Tocris Bioscience; X-tremeGENE™ HP DNA Transfection Reagent (6366244001) and Dithiothreitol (10197777001) from Roche.

## Generation of knockout, knockdown and cDNA overexpression cell lines

sgRNAs (oligonucleotide sequences are indicated below) were cloned into lentiCRISPR-v2 (Addgene) linearized with BsmBI and ligated by T4 ligase (NEB). sgRNA-expressing vector along with lentiviral packaging vectors Delta-VPR and CMV VSV-G were transfected into HEK293T cells using the XTremeGene 9 transfection reagent (Roche). Similarly, for overexpression cell lines, gBlocks (IDT) containing the cDNA of interest were cloned into pMXS-IRES-BLAST, pMXS-IRES-GFP and pMXS-IRES-PURO linearized with XhoI and NotI by Gibson Assembly (NEB). cDNA vectors along with retroviral packaging vectors gag-pol and CMV VSV-G were transfected into HEK293T cells. 48 hours after transfection, the virus-containing supernatant was collected and passed through a 0.45 μm filter to eliminate cells. Target cells in 6-well tissue culture plates were infected in media containing 4 µg/mL of polybrene and a spin infection was performed by centrifugation at 2,200 rpm for 1 hour. Post-infection, virus was removed, and cells were selected with puromycin (lentiCRISPR-v2, pMXS_IRES_Puro), blasticidin (pMXS_IRES_Blast) or by flow sorting of top 2% GFP positive cells (pMXS-IRES-GFP). Clones were validated for loss of the relevant protein via immunoblotting. *sgSLC33A1* transduced cells were validated through ICE analysis (Synthego) and immunoblotting.

### Oligo Sequences

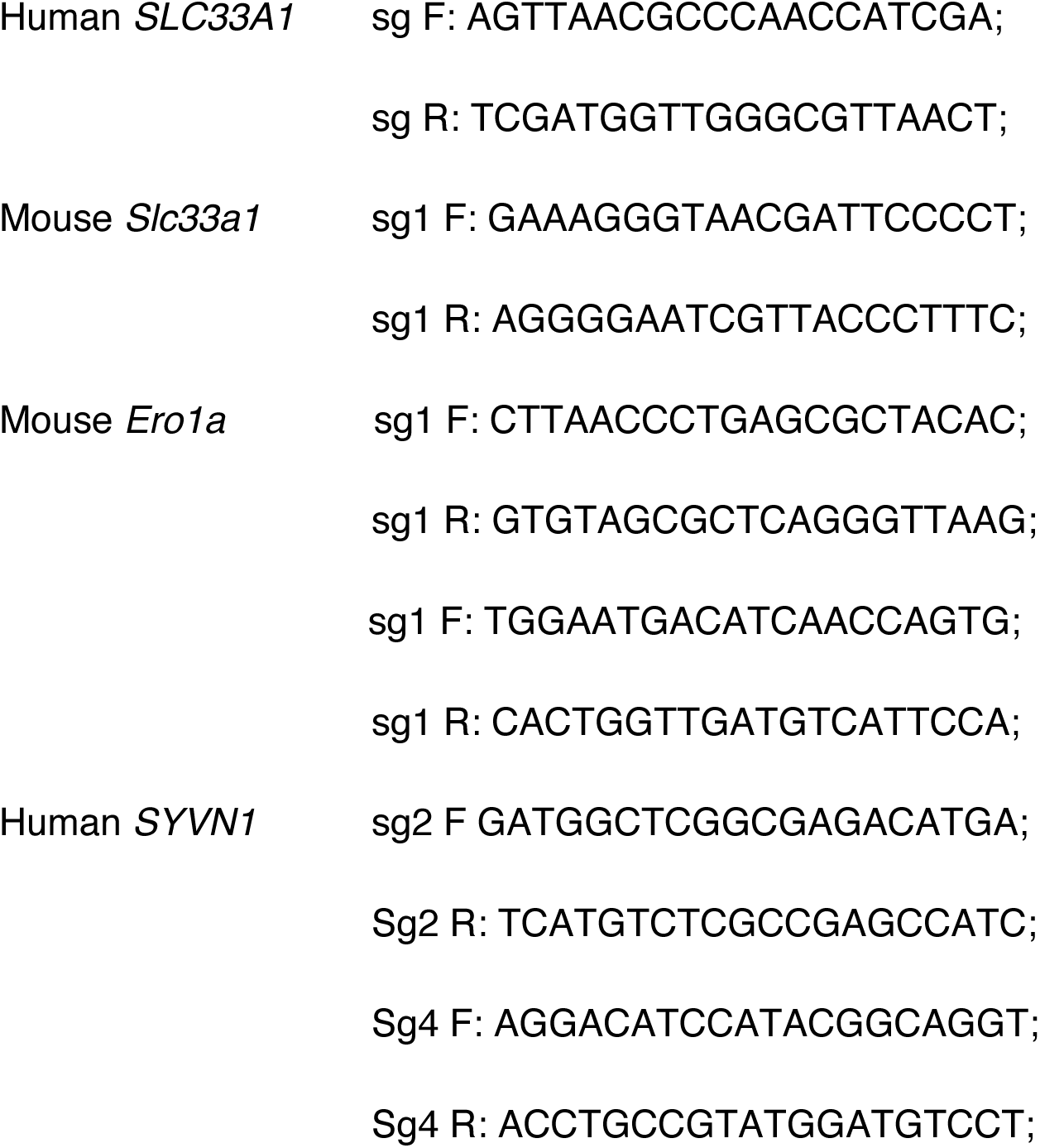

### EMC3-mScarlet-3xHA sequence

ATGGCAGGGCCAGAACTGTTGCTCGACTCCAACATCCGCCTCTGGGTGGTCCTACCCATCGTTATCATCACTTTCTTCGTAGGCATGATCCGCCACTACGTGTCCATCCTGCTGCAGAGCGACAAGAAGCTCACCCAGGAACAAGTATCTGACAGTCAAGTCCTAATTCGAAGCAGAGTCCTCAGGGAAAATGGAAAATACATTCCCAAACAGTCTTTCTTGACACGAAAATATTATTTCAACAACCCAGAGGATGGATTTTTCAAAAAAACTAAACGGAAGGTAGTGCCACCTTCTCCTATGACTGATCCTACTATGTTGACAGACATGATGAAAGGGAATGTAACAAATGTCCTCCCTATGATTCTTATTGGTGGATGGATCAACATGACATTCTCAGGCTTTGTCACAACCAAGGTCCCATTTCCACTGACCCTCCGTTTTAAGCCTATGTTACAGCAAGGAATCGAGCTACTCACATTAGATGCATCCTGGGTGAGTTCTGCATCCTGGTACTTCCTCAATGTATTTGGGCTTCGGAGCATTTACTCTCTGATTCTGGGCCAAGATAATGCCGCTGACCAATCACGAAT GATGCAGGAGCAGATGACGGGAGCAGCCATGGCCATGCCCGCAGACACAAACAAAGCTTTCAAGACAGAGTGGGAAGCTTTGGAGCTGACGGATCACCAGTGGGCACTAGATGATGTCGAAGAAGAGCTCATGGCCAAAGACCTCCACTTCGAAGGCATGTTCAAAAAGGAATTACAGACCTCTATTTTTACGCGTGGTGGTGGTTCTGGAGGCGGCGGATCAGTGAGCAAGGGCGAGGCAGTGATCAAGGAGTTCATGCGGTTCAAGGTGCACATGGAGGGCTCCATGAACGGCCACGAGTTCGAGATCGAGGGCGAGGGCGAGGGCCGCCCCTACGAGGGCACCCAGACCGCCAAGCTGAAGGTGACCAAGGGTGGCCCCCTGCCCTTCTCCTGGGACATCCTGTCCCCTCAGTTCATGTACGGCTCCAGGGCCTTCACCAAGCACCCCGCCGACATCCCCGACTACTATAAGCAGTCCTTCCCCGAGGGCTTCAAGTGGGAGCGCGTGATGAACTTCGAGGACGGCGGCGCCGTGACCGTGACCCAGGACACCTCCCTGGAGGACGGCACCCTGATCTACAAGGTGAAGCTCCGCGGCACCAACTTCCCTCCTGACGGCCCCGTAATGCAGAAGAAGACAATGGGCTGGGAAGCGTCCACCGAGCGGTTGTACCCCGAGGACGGCGTGCTGAAGGGCGACATTAAGATGGCCCTGCGCCTGAAGGACGGTGGCCGCTACCTGGCGGACTTCAAGACCACCTACAAGGCCAAGAAGCCCGTGCAGATGCCCGGCGCCTACAACGTCGACCGCAAGTTGGACATCACCTCCCACAACGAGGAATACACCGTGGTGGAACAGTACGAACGCTCCGAGGGCCGCCACTCCACCGGCGGCATGGACGAGCTGTACAAGGGCGGATCAGGTGGTGGTTCTTATCCCTATGACGTGCCTGATTACGCCGGCACAGGATCCTACCCCTATGATGTGCCTGACTACGCTGGCAGCGCCGGATACCCTTATGATGTGCCTGATTATGCTGAAGAGGATCTGTAA

### EMC3-mScarlet-3xMYC sequence

ATGGCAGGGCCAGAACTGTTGCTCGACTCCAACATCCGCCTCTGGGTGGTCCTACCCATCGTTATCATCACTTTCTTCGTAGGCATGATCCGCCACTACGTGTCCATCCTGCTGCAGAGCGACAAGAAGCTCACCCAGGAACAAGTATCTGACAGTCAAGTCCTAATTCGAAGCAGAGTCCTCAGGGAAAATGGAAAATACATTCCCAAACAGTCTTTCTTGACACGAAAATATTATTTCAACAACCCAGAGGATGGATTTTTCAAAAAAACTAAACGGAAGGTAGTGCCACCTTCTCCTATGACTGATCCTACTATGTTGACAGACATGATGAAAGGGAATGTAACAAATGTCCTCCCTATGATTCTTATTGGTGGATGGATCAACATGACATTCTCAGGCTTTGTCACAACCAAGGTCCCATTTCCACTGACCCTCCGTTTTAAGCCTATGTTACAGCAAGGAATCGAGCTACTCACATTAGATGCATCCTGGGTGAGTTCTGCATCCTGGTACTTCCTCAATGTATTTGGGCTTCGGAGCATTTACTCTCTGATTCTGGGCCAAGATAATGCCGCTGACCAATCACGAATGATGCAGGAGCAGATGACGGGAGCAGCCATGGCCATGCCCGCAGACACAAACAAAGCTTTCAAGACAGAGTGGGAAGCTTTGGAGCTGACGGATCACCAGTGGGCACTAGATGATGTCGAAGAAGAGCTCATGGCCAAAGACCTCCACTTCGAAGGCATGTTCAAAAAGGAATTACAGACCTCTATTTTTGGTGGTGGTTCTGGAGGCGGCGGATCAGTGAGCAAGGGCGAGGCAGTGATCAAGGAGTTCATGCGGTTCAAGGTGCACATGGAGGGCTCCATGAACGGCCACGAGTTCGAGATCGAGGGCGAGGGCGAGGGCCGCCCCTACGAGGGCACCCAGACCGCCAAGCTGAAGGTGACCAAGGGTGGCCCCCTGCCCTTCTCCTGGGACATCCTGTCCCCTCAGTTCATGTACGGCTCCAGGGCCTTCACCAAGCACCCCGCCGACATCCCCGACTACTATAAGCAGTCCTTCCCCGAGGGCTTCAAGTGGGAGCGCGTGATGAACTTCGAGGACGGCGGCGCCGTGACCGTGACCCAGGACACCTCCCTGGAGGACGGCACCCTGATCTACAAGGTGAAGCTCCGCGGCACCAACTTCCCTCCTGACGGCCCCGTAATGCAGAAGAAGACAATGGGCTGGGAAGCGTCCACCGAGCGGTTGTACCCCGAGGACGGCGTGCTGAAGGGCGACATTAAGATGGCCCTGCGCCTGAAGGACGGCGGCCGCTACCTGGCGGACTTCAAGACCACCTACAAGGCCAAGAAGCCCGTGCAGATGCCCGGCGCCTACAACGTCGACCGCAAGTTGGACATCACCTCCCACAACGAGGAATACACCGTGGTGGAACAGTACGAACGCTCCGAGGGCCGCCACTCCACCGGCGGCATGGACGAGCTGTACAAGGGAGGGAGCGGCGAGCAGAAGCTGATTTCTGAGGAAGATCTGGGCACAGGATC CGAACAGAAACTGATTTCTGAGGAAGATCTGGGCAGCGCCGGAGAGCAGAAGCTGATTTCTGAAGAGGATCTGTAA

### FLAG-*SLC33A1* sequence: FLAG tag

ATGGATTACAAGGATGACGATGACAAGGGCGGATCAGGTGGTGGTTCTTCACCCACCATCTCCCACAAGGACAGCAGCCGGCAACGGCGGCCAGGGAATTTCAGTCACTCTCTGGATATGAAGAGCGGTCCCCTGCCGCCAGGCGGTTGGGATGACAGTCATTTGGACTCAGCGGGCCGGGAAGGGGACAGAGAAGCTCTTCTGGGGGATACCGGCACTGGCGACTTCTTAAAAGCCCCACAGAGCTTCCGGGCCGAACTAAGCAGCATTTTGCTACTACTCTTTCTTTACGTGCTTCAGGGTATTCCCCTGGGCTTGGCGGGAAGCATCCCACTCATTTTGCAAAGCAAAAATGTTAGCTATACAGACCAAGCTTTCTTCAGTTTTGTCTTTTGGCCCTTCAGTCTCAAATTACTCTGGGCCCCGTTGGTTGATGCGGTCTACGTTAAGAACTTCGGTCGTCGCAAATCTTGGCTTGTCCCGACACAGTATATACTAGGACTCTTCATGATCTATTTATCCACTCAGGTGGACCGTTTGCTTGGGAATACCGATGACAGAACACCCGACGTGATTGCTCTCACTGTGGCGTTCTTTTTGTTTGAATTCTTGGCCGCCACTCAGGACATTGCAGTCGATGGTTGGGCGTTAACTATGTTATCCAGGGAAAATGTGGGTTATGCTTCTACTTGCAATTCGGTGGGCCAAACAGCGGGTTACTTTTTGGGCAATGTTTTGTTTTTGGCCCTTGAATCTGCCGACTTTTGTAACAAATATTTGCGGTTTCAGCCTCAACCCAGAGGAATCGTTACTCTTTCAGATTTCCTTTTTTTCTGGGGAACTGTATTTTTAATAACAACAACATTGGTTGCCCTTCTGAAAAAAGAAAACGAAGTATCAGTAGTAAAAGAAGAAACACAAGGGATCACAGATACTTACAAGCTGCTTTTTGCAATTATAAAAATGCCAGCAGTTCTGACATTTTGCCTTCTGATTCTAACTGCAAAGATTGGTTTTTCAGCAGCAGATGCTGTAACAGGACTGAAATTGGTAGAAGAGGGAGTACCCAAAGAACATTTAGCCTTATTGGCAGTTCCAATGGTTCCTTTGCAGATAATACTGCCTCTGATTATCAGCAAATACACTGCAGGTCCCCAGCCATTAAACACATTTTACAAAGCCATGCCCTACAGATTATTGCTTGGGTTAGAATATGCCCTACTGGTTTGGTGGACTCCTAAAGTAGAACATCAAGGGGGATTCCCTATATATTACTATATCGTAGTCCTGCTGAGTTATGCTTTACATCAGGTTACAGTGTACAGCATGTATGTTTCTATAATGGCTTTCAATGCAAAGGTTAGTGATCCACTTATTGGAGGAACATACATGACCCTTTTAAATACCGTGTCCAATCTGGGAGGAAACTGGCCTTCTACAGTAGCTCTTTGGCTTGTAGATCCCCTCACAGTAAAAGAGTGTGTAGGAGCATCAAACCAGAATTGTCGAACACCTGATGCTGTTGAGCTTTGCAAAAAACTGGGTGGCTCATGTGTTACAGCCCTGGATGGTTATTATGTGGAGTCCATTATTTGTGTTTTCATTGGATTTGGTTGGTGGTTCTTTCTTGGTCCAAAATTTAAAAAGTTACAGGATGAAGGATCATCTTCGTGGAAATGCAAAAGGAACAATTAA

Gene fragment of FLAG-SLC33A1 was purchased from Twist Biosciences and cloned directly into linearized pMXS-IRES-Blast (Cell Biolabs, RTV-016) or pMXS-IRES-GFP (Cell Biolabs, RTV-013).

### **ER-GshF-V5 sequence:** signal sequence (ERP44), V5 tag, KDEL retention signal

ATGCATCCTGCCGTCTTCCTATCCTTACCCGACCTCAGATGCTCCCTTCTGCTCCTGGTAACTTGGGTTTTTACTCCTGTAACAACTGAAGGCGGAAGCACCCTTAATCAGCTTCTCCAGAAGTTGGAGGCGACTTCCCCCATTCTCCAGGCGAACTTCGGGATAGAAAGGGAGTCATTGAGGGTTGACCGCCAGGGTCAGCTGGTCCACACACCGCACCCCTCATGTCTGGGAGCCCGCAGTTTTCATCCTTACATACAAACCGACTTTTGTGAATTCCAAATGGAACTGATTACACCAGTAGCCAAAAGTACGACGGAGGCCCGACGCTTTCTTGGCGCGATAACTGATGTAGCAGGACGAAGCATTGCAACTGACGAGGTGCTGTGGCCATTGAGTATGCCACCACGACTTAAAGCCGAGGAAATTCAAGTAGCGCAACTCGAGAACGACTTCGAAAGACATTATCGGAACTACTTGGCAGAGAAGTACGGCACCAAATTGCAGGCGATTAGTGGAATTCA TTACAATATGGAACTTGGGAAGGACTTGGTTGAGGCGCTTTTTCAAGAGTCAGATCAGACTGACATGATCGCATTTAAAAACGCTCTGTATCTCAAGCTCGCCCAGAACTATTTGAGGTATCGGTGGGTCATTACTTACCTGTTTGGAGCAAGTCCCATTGCAGAACAAGGATTCTTTGACCAAGAAGTGCCGGAGCCTATGCGCTCTTTCCGCAACTCCGACCACGGCTACGTTAACAAGGAAGAGATACAGGTAAGCTTTGTATCCCTTGAAGACTATGTCTCCGCGATCGAGACCTACATCGAGCAGGGTGACCTTATAGCCGAGAAAGAGTTTTACTCAGCCGTGCGCTTTAGAGGACAAAAAGTCAATCGCTCCTTCCTTGATAAGGGTATAACTTATCTGGAGTTCAGAAACTTTGACTTGAACCCATTTGAGAGAATAGGCATCAGTCAAACCACTATGGATACCGTTCACTTGCTCATACTGGCCTTTCTCTGGTTGGATAGTCCGGAAAACGTGGACCAGGCCCTTGCCCAGGGACACGCGCTTAACGAGAAAATAGCCCTCTCCCATCCATTGGAGCCCCTCCCTTCAGAGGCCAAGACACAGGACATCGTGACCGCACTCGACCAGCTCGTACAGCACTTCGGATTGGGAGATTACCACCAGGATCTCGTGAAACAAGTGAAAGCGGCGTTTGCTGATCCGAATCAAACCCTGTCAGCTCAACTTCTTCCTTATATTAAGGACAAGTCACTCGCAGAATTCGCTCTCAATAAGGCACTCGCATATCATGACTATGACTGGACCGCTCACTACGCCCTTAAAGGTTACGAAGAAATGGAGCTCAGTACGCAGATGCTGCTCTTTGATGCTATCCAGAAAGGAATACACTTCGAGATACTCGATGAGCAAGATCAGTTCTTGAAGCTGTGGCATCAAGATCACGTAGAATATGTTAAAAACGGTAATATGACCAGCAAGGATAACTATGTAGTACCTCTCGCAATGGCCAACAAGACTGTTACTAAGAAAATTCTTGCTGACGCTGGGTTCCCTGTTCCGTCCGGGGACGAATTTACTAGCTTGGAGGAGGGACTGGCCTACTACCCGCTTATTAAAGATAAGCAAATTGTAGTAAAGCCAAAGAGCACGAATTTCGGCTTGGGTATCAGCATCTTCCAAGAACCCGCCAGTCTCGACAATTATCAAAAAGCATTGGAAATAGCATTTGCGGAGGACACTAGTGTGCTCGTCGAAGAATTCATTCCAGGCACGGAATACCGATTCTTCATTTTGGACGGACGCTGTGAGGCAGTCCTTTTGAGGGTAGCTGCCAATGTAATAGGGGACGGGAAACACACAATCAGAGAGTTGGTAGCGCAGAAAAACGCAAATCCCCTGCGCGGTAGGGATCATAGATCACCCTTGGAAATCATAGAGCTTGGGGACATAGAGCAACTCATGCTGGCACAGCAGGGTTACACTCCAGATGACATCCTGCCAGAGGGTAAGAAAGTGAATTTGAGGCGGAACAGCAATATTAGTACTGGGGGAGACTCCATAGACGTCACAGAAACAATGGATAGCTCTTATCAAGAACTTGCAGCAGCGATGGCTACCAGTATGGGGGCATGGGCCTGTGGAGTTGATCTGATTATACCCGACGAAACGCAGATTGCCACAAAGGAAAATCCACATTGCACGTGTATTGAACTTAACTTCAACCCCTCCATGTACATGCATACATACTGCGCTGAGGGGCCGGGGCAGGCAATTACAACCAAAATACTCGACAAACTCTTCCCGGAGATCGTGGCCGGACAAACTGGAGGAAGCGGAGGAAGCGGAGGAAGCGATTACAAGGGTAAGCCTATCCCTAACCCTCTCCTCGGTCTCGATTCTACGAAGGATGAATTATAA

The Puro section of pCW57.1 (Addgene 41393) is replaced by GFP to generate pCW57.1-GFP. Gene fragment of ER-GshF was purchased from Twist Biosciences and cloned directly into linearized pCW57.1-GFP linearized by AgeI and NheI.

#### SLC33A1-GFP

ATGtcacccaccatctcccacaaggacagcagccggcaacggcggccagggaatttcagtcactctctggatatgaagagcggtcccctgccgccaggcggttgggatgacagtcatttggactcagcgggccgggaaggggacagagaagctcttctgggggataccggcactggcgacttcttaaaagccccacagagcttccgggccgaactaagcagcattttgctactactctttctttacgtgcttcagggtattcccctgggcttggcgggaagcatcccactcattttgcaaagcaaaaatgttagctatacagaccaagctttcttcagttttgtcttttggcccttcagtctcaaattactctgggccccgttggttgatgcggtctacgttaagaacttcggtcgtcgcaaatcttggcttgtcccgacacagtatatactaggactcttcatgatctatttatccactcaggtggaccgtttgcttgggaataccgatgacagaacacccgacgtgattgctctcactgtggcgttctttttgtttgaattcttggccgccactcaggacattgcagtcgatggttgggcgttaactatgttatccagggaaaatgtgggttatgcttctacttgcaattcggtgggccaaacagcgggttactttttgggcaatgttttgtttttggcccttgaatctgccgacttttgtaacaaatatttgcggtttcagcctcaacccagaggaatcgttactctttcagatttcctttttttctg gggaactgtatttttaataacaacaacattggttgcccttctgaaaaaagaaaacgaagtatcagtagtaaaagaagaaacacaagggatcacagatacttacaagctgctttttgcaattataaaaatgccagcagttctgacattttgccttctgattctaactgcaaagattggtttttcagcagcagatgctgtaacaggactgaaattggtagaagagggagtacccaaagaacatttagccttattggcagttccaatggttcctttgcagataatactgcctctgattatcagcaaatacactgcaggtccccagccattaaacacattttacaaagccatgccctacagattattgcttgggttagaatatgccctactggtttggtggactcctaaagtagaacatcaagggggattccctatatattactatatcgtagtcctgctgagttatgctttacatcaggttacagtgtacagcatgtatgtttctataatggctttcaatgcaaaggttagtgatccacttattggaggaacatacatgacccttttaaataccgtgtccaatctgggaggaaactggccttctacagtagctctttggcttgtagatcccctcacagtaaaagagtgtgtaggagcatcaaaccagaattgtcgaacacctgatgctgttgagctttgcaaaaaactgggtggctcatgtgttacagccctggatggttattatgtggagtccattatttgtgttttcattggatttggttggtggttctttcttggtccaaaatttaaaaagttacaggatgaaggatcatcttcgtggaaatgcaaaaggaacaatTCGAATTCTTTGGAAGTTTTGTTTCAAGGTCCAACTGCTGCCGCCGCTGTGAGCAAGGGCGAGGAGCTGTTCACCGGGGTGGTGCCCATCCTGGTCGAGCTGGACGGCGACGTAAACGGCCACAAGTTCAGCGTGTCCGGCGAGGGCGAGGGCGATGCCACCTACGGCAAGCTGACCCTGAAGTTCATCTGCACCACCGGCAAGCTGCCCGTGCCCTGGCCCACCCTCGTGACCACCCTGACCTACGGCGTGCAGTGCTTCAGCCGCTACCCCGACCACATGAAGCAGCACGACTTCTTCAAGTCCGCCATGCCCGAAGGCTACGTCCAGGAGCGCACCATCTTCTTCAAGGACGACGGCAACTACAAGACCCGCGCCGAGGTGAAGTTCGAGGGCGACACCCTGGTGAACCGCATCGAGCTGAAGGGCATCGACTTCAAGGAGGACGGCAACATCCTGGGGCACAAGCTGGAGTACAACTACAACAGCCACAACGTCTATATCATGGCCGACAAGCAGAAGAACGGCATCAAGGTGAACTTCAAGATCCGCCACAACATCGAGGACGGCAGCGTGCAGCTCGCCGACCACTACCAGCAGAACACCCCCATCGGCGACGGCCCCGTGCTGCTGCCCGACAACCACTACCTGAGCACCCAGTCCAAGCTGAGCAAAGACCCCAACGAGAAGCGCGATCACATGGTCCTGCTGGAGTTCGTGACCGCCGCCGGGATCACTCTCGGCATGGACGAGCTGTACAAGTCCGGAGGTGGTACTGAAACTTCTCAGGTTGCACCGGCTTAA

#### SLC33A1^MAP^ – GFP sequence

ATGtcacccaccatctcccacaaggacagcagccggcaacggcggccagggaatttcagtcactctctggatatgaagagcggtcccctgccgccaggcggttgggatgacagtcatttggactcagcgggccgggaaggggacagagaagctcttctgggggataccggcactggcgacttcttaaaagccccacagagcttccgggccgaactaagcagcattttgctactactctttctttacgtgcttcagggtattcccctgggcttggcgggaagcatcccactcattttgcaaagcaaaaatgttagctatacagaccaagctttcttcagttttgtcttttggcccttcagtctcaaattactctgggccccgttggttgatgcggtctacgttaagaacttcggtcgtcgcaaatcttggcttgtcccgacacagtatatactaggactcttcatgatctatttatccactcaggtggaccgtttgcttgggaataccgatgacagaacacccgacgtgattgctctcactgtggcgttctttttgtttgaattcttggccgccactcaggacattgcagtcgatggttgggcgttaactatgttatccagggaaaatgtgggttatgcttctacttgcaattcggtgggccaaacagcgggttactttttgggcaatgttttgtttttggcccttgaatctgccgacttttgtaacaaatatttgcggtttcagcctcaacccagaggaatcgttactctttcagatttcctttttttctggggaactgtatttttaataacaacaacattggttgcccttctgaaaaaagaaaacgaagtatcagtagtaaaagaagaaacacaagggatcacagatacttacaagctgctttttgcaattataaaaatgccagcagttctgacattttgccttctgattctaactgcaaagattggtttttcagcagcagatgctgtaacaggactgaaattggtagaagagggagtacccaaagaacatttagccttattggcagttccaatggttcctttgcagataatactgcctctgattatcagcaaatacactgcaggtccccagccattaaacacattttacaaagccatgccctacagattattgcttgggttagaatatgccctactggtttggtggactcctaaagtagaaGGGGATGGCATGGTCCCCCCCGGCcctatatattactatatcgtagtcctgctgagttatgctttacatcaggttacagtgtacagcatgtatgtttctataatggctttcaatgcaaaggttagtgatccacttattggaggaacatacatgacccttttaaataccgtgtccaatctgggaggaaactggccttctacagtagctctttggcttgtagatcccctcacagtaaaagagtgtgtaggagcatcaaaccagaattgtcgaacacctgatgctgttgagctttgcaaaaaactgggtggctcatgtgttacagccctggatggttattatgtggagtccattatttgtgttttcattggatttggttggtggttctttcttggtccaaaatttaaaaagttacaggatgaaggatcatcttcgtggaaatgcaaaaggaacaatTCGAATTCTTTGGAAGTTTTGTTTCAAGGTCCAACTGCTGCCGCCGCTGTGAGCAAGGGCGAGGAGCTGTT CACCGGGGTGGTGCCCATCCTGGTCGAGCTGGACGGCGACGTAAACGGCCACAAGTTCAGCGTGTCCGGCGAGGGCGAGGGCGATGCCACCTACGGCAAGCTGACCCTGAAGTTCATCTGCACCACCGGCAAGCTGCCCGTGCCCTGGCCCACCCTCGTGACCACCCTGACCTACGGCGTGCAGTGCTTCAGCCGCTACCCCGACCACATGAAGCAGCACGACTTCTTCAAGTCCGCCATGCCCGAAGGCTACGTCCAGGAGCGCACCATCTTCTTCAAGGACGACGGCAACTACAAGACCCGCGCCGAGGTGAAGTTCGAGGGCGACACCCTGGTGAACCGCATCGAGCTGAAGGGCATCGACTTCAAGGAGGACGGCAACATCCTGGGGCACAAGCTGGAGTACAACTACAACAGCCACAACGTCTATATCATGGCCGACAAGCAGAAGAACGGCATCAAGGTGAACTTCAAGATCCGCCACAACATCGAGGACGGCAGCGTGCAGCTCGCCGACCACTACCAGCAGAACACCCCCATCGGCGACGGCCCCGTGCTGCTGCCCGACAACCACTACCTGAGCACCCAGTCCAAGCTGAGCAAAGACCCCAACGAGAAGCGCGATCACATGGTCCTGCTGGAGTTCGTGACCGCCGCCGGGATCACTCTCGGCATGGACGAGCTGTACAAGTCCGGAGGTGGTACTGAAACTTCTCAGGTTGCACCGGCTTAA

#### Pmab-1 Fv-clasp

##### Heavy Chain

ATGAAGTATCTTCTCCCTACCGCCGCGGCGGGGTTGTTGTTACTTGCAGCTCAGCCGGCTATGGCACAAATTCAGCTGCAACAATCAGGAACCGTGTTAGTGAAACCTGCTTCCAGTGTCAAAATTTCATGCAAAGCATCCGGTTACTCGTTTACCTCGCACTACATGCACTGGATTAGACAGCAACCCGGACAAGGACTTGAGTGGATAGGATGGATAAGTCCTGAGCAAGGAAACACGAAGTATAACCAGAAGTTTGACGGGAAAGCTACGCTTACTGCAGACAAGAGTTCCAGCATCGCCTACATGCAATTATCCTCTTTAACTAGCGAAGACTCAGCGGTGTACTTTTGTGTTAGCTGGGAAGATTGGAGCGCATACTGGGGCCAGGGCACTCTTGTCACCGTGTGCTCCGGGAGCGACTACGAGTTCCTGAAAAGTTGGACCGTGGAAGACCTGCAAAAGCGTCTTTTGGCTTTAGATCCTATGATGGAGCAGGAGATCGAAGAGATTAGACAGAAATACCAATCTAAACGCCAACCCATCCTTGACGCCATCGAAGCAAAGCACCACCACCACCACCACTG A

##### Light Chain

ATGAAGTATCTTCTCCCTACCGCCGCGGCGGGGTTGTTGTTACTTGCAGCTCAGCCGGCTATGGCAGACATCGTGCTCACCCAATCTCCTGCGTTGACAGTCAGCCTGGGTCAACGTGCTACGATATCGTGTAAGACCAATCAAAATGTCGATTATTACGGCAACAGTTATGTGCACTGGTACCAGCAAAAGCCCGGACAAAAGCCAAAGCTTCTTATTTACCTGGCGTCAAATCTGGCGTCCGGCATCCCCGCACGCTTTTCCGGCCGGGGTAGTGGAACCGACTTCACGCTTACTATAGATCCGGTCGAGGCCGCTGATACCGCAACTTACTATTGCCAGCAATCACGCGACTTGCCTAACACGTTTGGTGCCGGGACTAAGTTAGAGTTAAAGCGCGGTTCTGACTACGAGTTCCTGAAGAGTTGGACCGTCGAAGATCTTCAGAAGCGTTTACTCGCTTTAGATCCCATGATGGAACAAGAGATCGAAGAGATCCGGCAGAAGTACCAATGTAAACGGCAACCGATCTTGGACGCTATCGAGGCCAAATAA

### List of other constructs in the manuscript

pMXS_IRES_Bleo_EMC3-Scarlet-3xHA; pMXS_IRES_Bleo_EMC3-Scarlet-3xMYC; pMXS_IRES_BLAST_FLAG-33A1; pMXS_IRES_GFP_FLAG-33A1; pLJC6-3XHA-TMEM192-Blast; pLJC6-FLAG-SLC33A1-Blast; pLJC6-FLAG-SLC33A1-Y225A-Blast; pLJC6-FLAG-SLC33A1-N229A-Blast; pLJC6-FLAG-SLC33A1-Q356A-Blast; pLJC6-FLAG-SLC33A1-Y390A-Blast; pLJC6-FLAG-SLC33A1-Y418A-Blast; pLJC6-FLAG-SLC33A1-Q422A-Blast; PCW57.1-ER-GshF-GFP, PCW57.1-cyto-GshF-GFP, lentiCRISPR-v2_sg_SLC33A1 (PURO); lentiCRISPR-v2_sg_Slc33a1 (PURO); lentiCRISPR-v2_sg2_ SYVN1 (PURO); lentiCRISPR-v2_sg4_ SYVN1 (PURO); lentiCRISPR-v2_sg1_ Ero1a (PURO); lentiCRISPR-v2_sg2_ Ero1a (PURO)

### Immunoblotting

Cells were washed with ice-cold PBS prior to lysis in triton lysis buffer (50 mM Tris–HCl, pH 7.5, 1 mM EDTA, 1% (v/v) Triton X-100, 150 mM NaCl) supplemented with protease inhibitor cocktail (EMD Millipore, 539134). Each lysate was sonicated, after centrifugation at 20,000 g at 4°C for 10 min, supernatants were collected. Sample protein concentrations were determined by using Pierce BCA Protein Assay Kit (Thermo Scientific) with bovine serum albumin as a protein standard. Samples were resolved on 10-20% SDS-PAGE gels (Invitrogen) and analyzed by standard immunoblotting protocol. Briefly, membranes were incubated with primary antibodies in 5% (w/v) nonfat milk in TBST at 4°C overnight with shaking. Membranes were washed three times in TBST. Membranes were incubated with secondary antibody for 1 hour at room temperature with shaking, including α-mouse IgG-HRP linked (Cell Signaling, 7076) and α-rabbit IgG-HRP linked (Cell Signaling, 7074), used at 1: 5000 dilution. Membranes were washed again three times in TBST before visualization with ECL substrate.

### AMS modification and immunoblotting

To obtain untreated, completely reduced or oxidized samples, cells were trypsinized and counted. 3e^6^ cells/sample were pretreated with either water, 10 mM DTT or 5 mM diamide in normal growth medium at 37°C for 5 min. Cells were then washed with PBS containing 25 mM NEM once and incubated in the same buffer for 20 min. Cells were washed with PBS to remove excessive NEM before lysed in lysis buffer (50 mM Tris–HCl, pH 7.5, 1 mM EDTA, 1% (v/v)Triton X-100, 1% (w/v) SDS, 150 mM NaCl) supplemented with protease inhibitor cocktail (EMD Millipore, 539134). Each lysate was sonicated, after centrifugation at 20,000 g at 4°C for 10 min, supernatants were collected and boiled at 98°C for 5 min to denature the proteins. After cooling down, samples were reduced by 10 mM Tris[2-carboxyethyl] phosphine (TCEP) for 20 min at room temperature and then alkylated by addition of 30 mM AMS for 1 hour at room temperature. Samples were prepared with non-reducing SDS sample buffer and resolved on 8% SDS-PAGE gels.

### CRISPR-based genetic screens

The ER-focused human sgRNA library was designed and screens performed as previously described^55^. Oligonucleotides for sgRNAs were synthesized by CustomArray and amplified by PCR and cloned into lentiCRISPRv2-Opti puro (Addgene 163126). In brief, the plasmid pool was used to generate a lentiviral library, which was transfected into HEK293T cells and used to generate viral supernatant. For ER-GshF screening, HEK293T cells expressing inducible ER-GshF were infected at a multiplicity of infection of 0.6 and selected with puromycin. An initial sample of 20 million cells was harvested and infected cells were cultured with or without 1 µg/ml doxycycline for approximately 14 population doublings. Final samples of 20 million cells were collected. For *SLC33A1* knockout screening, *SLC33A1* knockout HEK293T cells expressing a vector control or *SLC33A1* cDNA were infected at a multiplicity of infection of 0.6 and selected with puromycin. An initial sample of 20 million cells was harvested and infected cells were cultured for approximately 14 population doublings. Final samples of 20 million cells were collected. DNA was extracted (DNeasy Blood and Tissue kit, Qiagen). sgRNA inserts were PCR-amplified and barcoded by PCR, using primers unique to each condition. PCR amplicons were purified and sequenced (NextSeq 500, Illumina). Screens were analyzed using Python (v.2.7.13), R (v.3.3.2) and Unix (v.4.10.0-37-generic x86_64). The gene score for each gene was defined as the median log_2_ fold change in the abundance of each sgRNA targeting that gene. A complete list of differential gene scores for each screen is provided in Supplementary Table 7-8.

### Immunofluorescence microscopy

For immunofluorescence assays, 5,000 HEK293T cells were seeded onto coverslips coated with poly-d-lysine in 24 well plate. For ER-GshF localization, cells were seeded with 1 µg/ml doxycycline and cultured for 48 hours. For EMC3-3xHA localization, cells were cultured for 48 hours. The cells were fixed with 4% (v/v) paraformaldehyde in PBS at room temperature for 30 min. After three PBS washes, cells were permeabilized with 0.1% (v/v) Triton X-100 in PBS for 30 min. After three additional PBS washes, cells were blocked in 10% (w/v) BSA in PBS for 30 min. For EMC3-HA localization, the coverslips were incubated with anti-HA (CST, 2367S, 1: 200), anti-calnexin (CST, 2679S, 1:200) in 10% BSA in PBS at 4 °C for 16 hours. For ER-GshF localization, the coverslips were incubated with anti-V5 (Sigma, V8012-50UG, 1:500), anti-calnexin (CST, 2679S, 1:200) in 10% BSA in PBS at 4 °C for 16 hours. The coverslips were then washed 3 times with PBS and incubated with secondary (Alexa Fluor 568 anti-mouse and Alexa Fluor 488 anti-rabbit) 1:500 in 10% BSA in PBS for 1 hour at room temperature in the dark. Coverslips were then washed with PBS for 10 min and then incubated with a solution of 200 nM DAPI in the dark before being washed an additional 3 times with PBS. Finally, coverslips were mounted onto slides with Prolong Gold Antifade mounting medium (Invitrogen). Images were obtained with a Nikon A1R MP+ multiphoton confocal microscope (Nikon) using a 60×/1.4 DIC Plan-Apochromat oil immersion objective. Images were obtained with excitation wavelengths as follows: DAPI 425–475 nm, Alexa Fluor 488 500–550 nm, Alexa Fluor 568 570–620 nm.

### Cystine isotope labeling

HEK293T cells expressing a vector control or inducible ER-GshF (iER-GshF) were pretreated with or without 1 mM BSO and 1 µg/ml doxycycline in dialyzed FBS for 24 hours before switching to cystine-labeled media with the same pretreatments and labeled for 8 hours before harvesting. Cystine labeled media were made by dissolving RPMI 1640 medium deficient w/o glutamine, methionine, cystine (Powder, from USBiological life sciences) supplemented with 4 mM glutamine, 0.2 mM methionine and 0.2 mM isotope labeled cystine (³C₆, ¹⁵N₂), and 10 % dialyzed FBS and 1% penicillin and streptomycin. After 8 hour incubation, the cells were harvested for ER-IP and metabolomic analysis.

### Cell proliferation assays

For cell proliferation assays with cell counting (ER-GshF and sg*SYVN1 assays*), cells were plated in triplicates in 6-well plates with at 10,000 cells per well, under the conditions described in each experiment. Cells were collected after 3-4 days and counted by Z2 Coulter Counter (Beckman). For cell proliferation assays with Cell Titer Glo measurement (tunicamycin and thapsigargin assays), cells were cultured in quintuplicate in 96-well plates at 1,000 cells per well in 200 µl DMEM medium under the conditions described in each experiment. after 4-5 days (with varying treatment conditions), 40 μl of Cell Titer Glo reagent (Promega) was added to each well, mixed briefly, and the luminescence read on a luminometer (Molecular Devices). Cell culture images were taken using a Zeiss Primovert equipped with Axio 503 mono camera and Zen 2 blue edition 2011 software.

### SLC33A1 coevolution analysis

The PhyloGene web server was queried to identify the top 200 genes with protein conservation and evolutionary trends across eukaryotes concordant with that of SLC33A1. Single-cell data from otherwise healthy mice across a range of ages was retrieved from the Tabula Muris Senis initiative. Normalized h5ad files from liver, pancreas, nonmyeloid brain cells, and myeloid brain cells were loaded into R and aggregated by taking the average expression of all cells at each timepoint and filtering out all genes with zero counts at either early timepoints (1mo and 3mo) or late timepoints (18mo, 21mo, 24mo, and 30mo). The Pearson correlation between the expression of SLC33A1 and each other gene in the dataset was computed to identify genes that both co-evolve with SLC33A1 and follow a similar pattern of expression as SLC33A1 over the course of the murine lifespan. For the purposes of visualization, genes with a coevolutionary Pearson correlation of at least 0.75 were defined as highly co-evolving.

### Rapid ER purification for metabolite profiling

ER were purified from HEK293T, HeLa, KP1 or KPK cells expressing EMC3-mScarlet-3xHA (ER-tag) or EMC3-mScarlet-3xMYC (control-tag). In brief, cells cultured in 2 x 15 cm dish were collected and washed twice with cold saline (0.9% NaCl), scraped into 3 ml of cold KPBS, and pelleted via centrifugation at 1,500 g at 4°C for 2 min. Cells were resuspended in 1 ml KPBS, 10 µl of cells were transferred into 90 µl of 1% (v/v) triton lysis buffer supplemented with protease inhibitor cocktail for a whole-cell protein sample and 10 µl of cells were transferred into 90 ml of ACN:MeOH:H_2_O (2:2:1), containing heavy labeled amino acid standards (Cambridge Isotope Labs, MSK-A2-1.2), for direct extraction of whole cell metabolites. With one set of 30 strokes and another set of 30 strokes, the remaining sample was homogenized using a 2-ml homogenizer. After centrifugation at 3,000 g for 10 min at 4 °C, the homogenate was incubated with 300 ml of KPBS pre-washed anti-HA magnetic beads (Thermo Scientific Pierce 88837) on a rotator shaker for 5 min at 4 °C. Beads were washed three times in cold KPBS, then 10% of bead volume was lysed with 50 µl 1% triton buffer for protein extracts and the remaining 90% was extracted in 50 µl ACN:MeOH:H_2_O (2:2:1) containing heavy labeled amino acid standards by vortex for 10 seconds followed by a rotator shaker for 10 min at 4 °C. After magnetic removing the beads, samples were spun down at 20,000g for 15 min to remove potential cellular debris or bead contamination. Samples were subjected to LC-MS polar metabolite profiling without drying. Data were normalized to protein level (BCA) or NAD^+^ abundance for whole cell metabolites, or UDP-GlcNAc/GalNAc for ER metabolites (see details in figure legends).

### LC/MS analysis for polar metabolites

Ultra-high performance liquid chromatography coupled with mass spectrometry (UHPLC/MS) analyses were conducted using a Thermo Scientific Vanquish Flex UHPLC system, interfaced with a Thermo Scientific Orbitrap ID-X Mass Spectrometer. For the separation of polar metabolites, a HILICON iHILIC-(P) Classic HILIC column (100 x 2.1 mm, 5 µm) with a HILICON iHILIC-(P) Classic guard column (20 x 2.1 mm, 5 µm) was utilized. The mobile-phase solvents consisted of solvent A = 20 mM ammonium bicarbonate, 2.5 µM medronic acid, 0.1% ammonium hydroxide in 95:5 water: acetonitrile and solvent B = 95:5 acetonitrile: water. The column compartment temperature was maintained at 45°C, and metabolites were eluted using a linear gradient at a flow rate of 250 µl/min as follows: 0-1 min, 90% B; 12 min, 35% B; 12.5-14.5 min, 25% B; 15 min, back to 90% B. The injection volume was 4 µl. Data was acquired in both positive and negative ion modes with the following settings: spray voltage, 3.5 kV/-2.8 kV; sheath gas, 50; auxiliary gas, 10; sweep gas, 1; ion transfer tube temperature, 300°C; vaporizer temperature, 200°C; mass range, 67-1000 Da; resolution, 120,000 (MS1), 30,000 (MS/MS); maximum injection time, 200 ms (MS1), 100 ms (MS/MS); isolation window, 1.5 Da. The LC/MS data were then processed and analyzed using XCMS, CompoundDiscoverer, and Skyline^56^.

### Whole-cell and ER proteomics

#### Sample preparation

ER were purified from HEK293T expressing EMC3-mScarlet-3xHA (ER-tag) or EMC3-mScarlet-3xMYC (control-tag) as described above. For whole-cell protein samples, after resuspension of the cell pellet in 1 ml KPBS, 10 μl of cells were transferred into 90 µl of 1% triton lysis buffer (50 mM Tris–HCl, pH 7.5, 1 mM EDTA, 1% (v/v) Triton X-100, 150 mM NaCl) supplemented with protease inhibitor cocktail (EMD Millipore, 539134). For ER-IP samples, following homogenization and centrifugation at 3,000 g for 10 min at 4 °C, the supernatants were passed through a 27G syringe needle five times prior to incubation with HA magnetic beads. After washed three times in cold KPBS, beads were resuspended in 100 µl of 1% triton lysis buffer supplemented with protease inhibitor cocktail.

#### Digestion

Samples were dried and dissolved in 8 M urea, 50 mM triethylammonium bicarbonate (TEAB) and 10 mM dithiothreitol (DTT), and disulfide bonds were reduced for 1 hour at room temperature. Alkylation was performed using iodoacetamide (IAA) for 1 hour at room temperature in the dark. Proteins were precipitated by Wessel/Flügge extraction and pellets were dissolved in 100 mM TEAB with endopeptidase LysC (2% w/w, enzyme/substrate) and incubated at 37 °C for 2-3 hours. Sequencing-grade modified trypsin (2 % w/w, enzyme/substrate) was added, and digestion proceeded overnight.

#### Labelling and fractionation

Peptide solutions were labelled with 270 µg aliquots of TMTpro (Thermo Scientific) for 1 h at room temperature and subsequently quenched with hydroxylamine for 15 min. An aliquot from each sample was combined for a ratio check, according to which the samples were mixed. The pooled sample was purified using a high-capacity reverse phase cartridge (Oasis HLB, Waters) and the eluate was fractionated using high pH reverse phase spin columns (Pierce) according to manufacturer specifications, yielding eight fractions.

#### LC-MS/MS

Fractionated peptides were analyzed using an Easy-nLC 1200 HPLC equipped with a 250 mm × 75 µm Easyspray column connected to a Fusion Lumos mass spectrometer (all Thermo Scientific) operating in synchronous precursor selection (SPS)-MS3 mode34 (10 SPS events). Solvent A was 0.1% formic acid in water and solvent B was 80% acetonitrile, 0.1% formic acid in water. Peptides from the ER immunoprecipitation were separated across a 90-min linear gradient and peptides from the whole-cell lysate were separated across a 120-min linear gradient going from 7 to 33% solvent B at 300 nl min−1. Precursors were fragmented by CID (35% CE) and MS2 ions were measured in the ion trap. MS2 ions were fragmented by HCD (65% CE) and MS3 reporter ions were measured in the orbitrap at 50K resolution. Data were analyzed using Skyline v.20.1.1.158.

#### Data analysis

Raw files were searched through Proteome Discoverer v.2.3 (Thermo Scientific) and spectra were queried against the human proteome using Sequest HT with a 1 % false discovery rate (FDR) applied. Oxidation of M was applied as a variable modification and carbamylation of C was applied as a static modification. A maximum isolation interference of 50% was allowed and 80% matching SPS ions were required. Protein abundance values were used for further statistical analysis. A complete list of identified peptides is provided in Supplementary Table 1-2.

Subsequent statistical analysis was performed within the Perseus framework. All values were log_2_-transformed and normalized to the median intensity within each sample. An FDR-corrected t-test (adjusted p-value = 0.05) was used to test for significant differences between sample groups. Imputed protein intensity values and precalculated p-values were loaded into R for analysis and subcellular localizations were generated as described below. Volcano plots were produced by filtering to proteins with at least 20 peptide spectrum matches, with proteins defined as hits if they passed a log-fold change cutoff of 2.5 and adjusted p-value cutoff of 0.05.

#### Annotation of subcellular localization

To associate proteins with the subcellular localization, the Cellular Compartment annotations from the Gene Ontology database were retrieved from BiomaRt for all genes in each RNAseq or proteomics dataset. Organelle-level annotations were generated from these granular classifications by searching for cellular compartment names containing unique minimal substrings (for instance, “endoplasmic| ER” for endoplasmic localization or “nucleus|nuclear” for nuclear localization). The list of ER proteins generated by this automated approach was supplemented by a previously identified list of ER proteins integrated from recently published studies. These classifications were used for downstream plotting and analysis.

### ER purification for RNA-Sequencing

#### Whole cell and ER RNA extraction

For RNA extraction, ER-IP was performed in the presence of RNAese inhibitor (Promega, N2111, 1:400). For whole cell RNA extraction, cell pellets collected before ER purification were lysed with 1 ml TRIzol (Fisher Scientific, 15-596-026); For ER RNA extraction, at the final step after the third KPBS wash, the entire bead volume was lysed in 1 ml TRIzol. 200 µl of chloroform was added and the mixture was incubated at room temperature for 2 min. After centrifugation 20,000 g at 4°C for 15 min, the upper phase was transferred into a new tube. 20 µl of GlycoBlueand and 500 µl ice-cold isopropanol were added. After mixing, the mixture was incubated at –20 °C for 20 min to precipitate the RNA. After centrifugation 20,000 g at 4°C for 15 min, the supernatant was discarded, and the pellet was washed with 1ml of 70% EtOH in DEPC water. The supernatant was then discarded, and the pellet was air dried and resuspended in 50 µl DEPC water. RNA was stored in –80°C before analysis.

#### RNA-Seq data analysis

Demultiplexed fastqs were pseudoaligned with Salmon to mm10 in the case of mouse samples or hg38 in the case of human samples. The resulting .sf files were loaded into R with tximport and normalized by limma-voom. In volcano plots, genes were defined to be hits if they passed a log-fold change cutoff of 0.5 and adjusted p-value cutoff of 0.05. Genes were defined to be highly and significantly enriched in one condition over another if and only if they passed a log-fold change cutoff of 2 and adjusted p-value cutoff of 0.01. A complete list of mRNA is provided in Supplementary Table 3 and 11.

### ER purification for lipidomic profiling

#### Sample preparation

ER were purified from HEK293T expressing EMC3-mScarlet-3xHA (ER-tag) or EMC3-mScarlet-3xMYC (control-tag) as described above. For whole-cell protein samples, after resuspension of the cell pellet in 1 ml KPBS, 10 μl of cells were transferred into 100 uL of PBS with 0.01% (w/v) of butylated hydroxytoluene to prevent lipid oxidation. Samples were stored at –80 °C before extraction. For ER-IP samples, after washed three times in cold KPBS, beads were resuspended in 100 µl of PBS with 0.01% (w/v) of butylated hydroxytoluene and stored at –80 °C before extraction.

#### Lipid extraction

To extract the lipids, samples were thawed on ice, 2 µl of SPLASH®LIPIDOMIX® (Avanti Polar Lipids, Inc Alabaster, AL, USA) and 2 µl Cer/Sph Mixture I ((Avanti Polar Lipids Inc) were added, and samples incubated for 15 min on ice. Next, 375 µl methanol (ULC-MS grade, > 99.97%; Biosolve B.V.), vortexed, followed by addition of 750 µl chloroform (EMSURE ACS, ISO, Reag. Ph Eur; Supleco). Samples were incubated under rotation (40 rpm) at 4°C for 1 hour. For phase separation 625 µl water were added, vortexed and samples incubated further 10 min as above. Samples were centrifuged for 10 min at 10.000 g 4°C and the lower organic phase was collected while tubes were placed in a magnetic rack. Organic (lipid containing) phases were dried under vacuum (40 mbar, 1200 rpm) and stored at –80°C until MS analysis.

#### LC-MS/MS analysis for lipids

Lipid extracts were reconstituted in 50 µl 2-propanol (ULC/MS-CC/SFC grade, >99.95%; Biosolve B.V), shaken for 15 min, centrifuged at 10.000 g for 10 min, and 40 µl of each individual sample transferred to a glass vial with a micro glass insert. Lipids were separated by reverse phase chromatography (Accucore C30 column; 150 mm x 2.1 mm 2.6 µM 150 Å, Thermo Fisher Scientific) using a Vanquish Horizon UHPLC system (Thermo Fisher Scientific, Germering, Germany) coupled on-line to an Orbitrap Exploris 240 mass spectrometer (Thermo Fisher Scientific, Bremen, Germany) equipped with a HESI source. Lipids were separated at a flow rate of 0.3 mL/min and a column temperature of 50°C by a gradient elution starting at 10% B increasing in 20 min to 80% B, followed by an increase to 95% B within 4 min, and further 3 min until 100% B is reached and kept for 5 min, followed by a decrease to 10%B within 0.1 min and equilibration at 10% B for 7.9 min^57^. Eluent A consisted of acetonitrile:water (50:50, v/v, both ULC/MS-CC/SFC grade, Biosolve B.V.) and Eluent B of 2-propanol:acetonitrile:water (85:10:5, v/v/v), both containing 5 mM ammonium formate (MS grade, Sigma Aldrich) and 0.1% (v/v) formic acid (ULC/MS-CC/SFC grade, Biosolve B.V.).

Full MS scans were acquired in positive and negative ionization mode with the following settings: spray voltage 3500 V/– 2500 V, sheath gas – 40 arb units, aux gas – 10 arb units, sweep gas – 1 arb unit, ion transfer tube – 300°C, vaporizer temperature – 370°C, EASY-IC run-start, default charge state – 1, resolution at m/z 200 – 120.000, normalized AGC target – 100%, maximum injection time – 100 ms, RF lens – 35%. For positive mode a scan range from m/z 200-1000 was covered and in negative mode full scans were acquired from m/z 200 – 1000 (until min 20) and from m/z 600-1700 (min 20 to 40).

Data-dependent acquisition (DDA) scans were acquired based on a cycle time of 1.3 s, a resolution of 15.000 at m/z 200, isolation window – 1.2 m/z, normalized stepped collision energies –17, 27, 37%, AGC target – 100%, maximum injection time – 60 ms, dynamic exclusion for 6 s after two-times fragmentation within 6 s.

#### Data analysis for lipids

Acquired raw files were processed in Lipostar2 (Version 2.1.5b1). Briefly, super sample filters retaining only lipids with isotopic patterns and MS/MS were applied and remaining MS/MS spectra searched against LIPID MAPS structural database (LMSD, downloaded 21.10.2023; addition of lipid standards using LipoStar DB manager). For identification, lipids with 3 and 4 stars were automatically approved, and searched for matching adducts and in-source fragments. Adducts were clustered and only approved features remained. These proposed identifications were manually checked for fragment matches and peak integration. Kendrick mass defect plots were used to validate the obtained identifications after manual inspection to filter out further false positive identifications. The obtained data matrix was exported and quantification performed in Microsoft Excel 2016 by dividing the peak area of the identified lipid by the peak area of the corresponding internal standard, multiplied by the concentration of the internal standard. Total pmol of each lipid were further normalized to the total protein amount obtained from BCA assay [pmol/µg]. Finally, these values were corrected by a normalization factor to consider the initial protein content of each sample prior to organelle isolation. Data were further filtered by keeping lipids quantified at least in 75% of the analyzed replicates per condition. ER and mitochondria specific lipids having at least a fold-change of 5 relative to the control-IP were selected. For each obtained lipidome (ER, mitochondria, whole cell), mol percent were calculated and used for final evaluation.

### [^3^H] GSSG uptake assay in immunopurified ER

*SLC33A1* knockout HEK293T or *Slc33a1* knockout KPK cells expressing EMC3-mScarlet-3xHA (ER-tag) complemented with a vector control or *SLC33A1* cDNA were treated with 1 mM BSO for 48 hours prior for ER isolation. ER were purified as described above. After three washes with KPBS using an electronic multi-channel pipette (Thermo Scientific, 4672100BT), beads were pooled and then evenly redistributed into tubes for different time points or GSSG concentrations. Uptake reactions were initiated by adding the indicated concentrations of unlabeled and radiolabeled GSSG (as specified in the figure legends) using the multi-channel pipette to ensure simultaneous reaction initiation across replicates. After the indicated incubation times, reactions were quenched by adding 1 mL cold KPBS, followed by two additional washes in 1 mL cold KPBS using the multi-channel pipette. Following the third wash, beads were extracted in ACN:MeOH:H_2_O (2:2:1) by vortex for 10 min. After removing the magnetic beads, samples were spun down at 20,000 g for 10 min to remove the remaining beads, and the supernatant was transferred to a scintillation vial with 5 mL of Insta-Gel Plus scintillation cocktail (Perkin Elmer #601339). Radioactivity was measured with the TopCount scintillation counter (Perkin Elmer).

### [^3^H] GSH uptake assay in immunopurified ER

*SLC33A1* knockout HEK293T expressing EMC3-mScarlet-3xHA (ER-tag) complemented with a vector control or *SLC33A1* cDNA were treated with 1 mM BSO for 48 hours prior for ER isolation. ER were purified as described above. After three times of KPBS wash, beads were incubated with 400 µM GSH containing 4 µM [^3^H] GSH for the indicated time. For GSSG competition assay, after three times of KPBS wash, beads were incubated with 80 µM unlabeled GSSG and 10 µM [^3^H]GSSG in the presence or absence of 5 mM unlabeled GSSG for 40 minutes. Uptake was stopped with the addition of 1 ml cold KPBS, and beads were subsequently washed two more times in 1 ml cold KPBS. Following the third wash, beads were extracted in ACN:MeOH:H_2_O (2:2:1) by vortex for 10 min. After magnetic removing the beads, samples were spun down at 20,000 g for 10 min to remove the remaining beads, and the supernatant was transferred to a scintillation vial with 5 mL of Insta-Gel Plus scintillation cocktail (Perkin Elmer #601339). Radioactivity was measured with the TopCount scintillation counter (Perkin Elmer).

### M+2 acetyl-CoA uptake assay in immunopurified ER

ER were purified as described above from *SLC33A1* knockout HEK293T expressing EMC3-mScarlet-3xHA (ER-tag) complemented with a vector control or *SLC33A1* cDNA. After three times of KPBS wash, beads were incubated with 400 µM m+2 acetyl-CoA for the indicated time. Uptake was stopped with the addition of 1 ml cold KPBS, and beads were subsequently washed two more times in 1 ml cold KPBS. Following the third wash, beads were extracted in ACN:MeOH:H_2_O (2:2:1) by vortex for 10 min. After magnetic removing the beads, samples were spun down at 20,000 g for 10 min to remove the remaining beads. Samples were subjected to LC-MS polar metabolite profiling without drying.

### Oxidized cysteine reactivity profiling

#### Sample preparation

*Slc33a1* knockout KPK cells expressing a vector control or *SLC33A1* cDNA were trypsinized and counted. 3e^6^ cells/ sample were washed with PBS and incubated twice in PBS containing 25 mM NEM for 10 minutes. Cells were washed with PBS to remove excessive NEM before lysing in triton lysis buffer (50 mM Tris–HCl, pH 7.5, 1 mM EDTA, 1% (v/v) Triton X-100, 150 mM NaCl) supplemented with protease inhibitor cocktail (EMD Millipore, 539134). Each lysate was sonicated by probe sonicator for 5 s at amplitude of 50 on ice. Following centrifugation (20,000 x g at 4 °C for 10 min), supernatants were collected, and sample protein concentrations were determined using Pierce BCA Protein Assay Kit (Thermo Scientific, 23225) with bovine serum albumin as a protein standard. Protein concentrations were adjusted to 2 mg/mL and samples (100 µl, 200 µg total proteins/sample) were added to new eppendorf tubes containing pre-weighed urea (48 mg/sample). Iodoacetamide (5 µl, 400 mM fresh stock in water, final concentration: 20 mM) was added and samples were incubated 37 °C for 1 hour. Proteins were then precipitated by addition of 500 µl methanol (pre-cooled at –80 °C), 400 µl LCMS-grade water, and 100 µl chloroform, and the mixture was vortexed and centrifuged (10,000 g, 10 min, 4°C) to afford a protein disc at the interface of CHCl3 and aqueous layers. The top layer was aspirated without perturbing the disk, additional MeOH (400 µl) was added, and the proteins were pelleted (10,000 g, 10 min, 4°C). The pellets were resuspended by addition of 90 µl of buffer containing urea (8 M), TCEP (5 mM) and Triethylammonium bicarbonate (TEAB, 50 mM, pH 8.5) with sonication. Following incubation at 56°C for 30 min, IA-DTB (10 µl 200 mM fresh stock in DMSO, final concentration: 20 mM) was added and samples were incubated 37 °C for 1 hour with shaking. Proteins were then precipitated by addition of 500 µl methanol (pre-cooled at –80 °C), 400 µl LCMS-grade water, and 100 µl chloroform. and the mixture was vortexed and centrifuged (10,000 g, 10 min, 4°C) to afford a protein disc at the interface of CHCl_3_ and aqueous layers. The top layer was aspirated without perturbing the disk, additional MeOH (400 μl) was added, and the proteins were pelleted (10,000 g, 10 min, 4°C) and the pellet were resuspended in 300 µL of 50 mM TEAB, containing 2 M urea by water bath sonication.

#### Trypsin/LysC digestion and streptavidin enrichment

Samples were digested by adding 3 µl of 100 mM CaCl_2_ (final concentration: 1 mM) and 2 µg of trypsin/LysC (20 µg trypsin/LysC resuspended in 40 µl trypsin/LysC buffer, then adding 4 µl to each sample), followed by shaking at 37 °C for 16 hours. Following centrifugation (20,000 x g, 5 min top remove undigested material), 250 µl of each sample was transferred into a new tube containing 250 µl of 1^st^ wash buffer (50 mM TEAB, 150 mM NaCl, 0.2% NP-40) containing 50 µl of streptavidin agarose beads that were pre-washed twice by the 1^st^ wash buffer). The samples were rotated at room temperature for 3 hours before washing with the 2^nd^ wash buffer (50 mM TEAB, 150 mM NaCl, 0.1% NP-40) 3 times, 3 times with PBS and 3 times with water. Peptides were eluted by addition of 300 µl of buffer containing 50% HPLC grade acetonitrile, 50% HPLC grade H_2_O, 0.1% FA and the eluate was then evaporated to dryness in Speed vac. Peptides were resuspended in 100 µl 30% ACN in 200 mM EPPS (pH 8.0) by water bath sonication for 5 minutes.

#### TMT labeling

Each sample was labeled by 3 µl of the corresponding 6-plex TMT tag (20 µg/µl) at room temperature for 1 h. 3 µl of 5% hydroxylamine was added into each sample and incubated at room temperature for 15 minutes to quench the unreacted tags. 5 µl of formic acid was then added into each sample before combining the samples together into a new tube. Samples were then evaporated to dryness in Speed vac and resuspended in 500 µl Velos buffer A (95% H_2_O, 5% CH_3_CN, 0.1% FA) with additional 20 µl of 20% FA to ensure the samples were acidic. Samples were desalted by using Sep-Pak C18 Cartridge and eluted by 1 mL Velos buffer B (80% CH_3_CN /0.1% FA). Samples were then evaporated to dryness in Speed vac and resuspended in 500 µl Buffer A (95% H_2_O, 5% CH_3_CN, 0.1% FA) with sonication and loaded on HPLC for offline high pH fractionation following conditions adapted from previous work^58^.

#### TMT liquid chromatography mass-spectrometry

Samples were analyzed by liquid chromatography tandem mass spectrometry. Cysteine reactivity samples were analyzed without the Real-Time Search feature while unenriched proteomics were analyzed separately with the Real-Time Search feature. The MS1, MS2, and MS3 files were extracted from RAW files using RawConverter (v1.2.0.1, available at https://github.com/proteomicsyates/RawConverter), uploaded to Integrated Proteomics Pipeline (IP2) and searched using the ProLuCID algorithm with a reverse concatenated version of the Mouse UniProt database (release-2024-03). For the cysteine reactivity dataset, peptides were searched with a static modification for carboxyamidomethylation (+57.02146 Da) and a differential modification for the IA-DTB probe (+239.1634 Da). Static modifications for *N*-termini and lysine residues corresponding to the 6-plex TMT tag (+229.1629 Da) were set. For the unenriched proteomics dataset, peptides were searched with a static modification for carboxyamidomethylation (+57.02146 Da) and a differential modification for oxidation of methionine (+15.9949 Da). Static modifications for *N*-termini and lysine residues corresponding to the 16-plex TMT tag (+304.2071 Da) were set. For both cysteine reactivity and unenriched proteomics, peptides were required to have at least one tryptic terminus and be at least 5 amino acids long. To achieve a peptide false-positive rate below 1%, ProLuCID data was filtered through DTASelect. The MS3-based peptide quantification was performed with reporter ion mass tolerance set to 20 ppm with IP2.

### Data processing and analysis for oxidized cysteine reactivity profiling and whole cell unenriched proteomics

Three 6-plex cysteine reactivity experiments with *Slc33a1* knockout KPK cells were analyzed (2 experimental conditions in triplicate for each donor). One 8-plex unenriched proteomics experiment with *Slc33a1* knockout KPK cells was analyzed (4 experimental conditions in triplicate with 4 empty channels). The following filters were used to process each experiment individually: removal of non-unique peptides, removal of half-tryptic peptides, removal of peptides with more than 2 internal missed cleavage sites, removal of peptides with low (<5,000) average of reporter ion intensities across all technical replicate groups, and removal of peptides with high variation between all technical replicate groups (coefficient of variation > 0.5). For cysteine reactivity profiling experiments, peptide ratios (signal intensity/sum of signal intensities per peptide) were calculated and cysteine aggregation was performed to aggregate signal intensities for multiple cysteines on the same peptide and multiple peptides for the same cysteine. Peptides were required to be quantified in a minimum of 2 experiments to be included in the final table. Experiments were combined and replicates were averaged (median) per condition. For unenriched proteomics experiment, peptide ratios were calculated and ratios of all peptides per protein were averaged. Dot plots and boxplots were created using R (v4.1.1), the ggplot2 (v3.5.1) and ggbeeswarm (v0.7.2) packages. The processed data can be found in Supplementary table 9-10.

### Protein expression and purification

#### SLC33A1 and SLC33A1^MAP^

Genes encoding human SLC33A1 and SLC33A1^MAP^ were subcloned into a pBacMam expression vector with C-terminal eGFP tag fused through a short linker containing a PreScission protease site^59^. The plasmid was mixed 1:3 (w/w) with PEI MAX**®** 40,000 (Polysciences, 24765-1) for 20 min at room temperature and then used to transfect Expi293 cells at 37°C in 8% CO2 (Thermo A14527). For a 1-L culture, 1 mg plasmid and 3 mg PEI MAX**®** were used. After 24 h incubation at 37 °C, valproic acid sodium salt (Sigma-Aldrich, P4543) was added to the culture at a final concentration of 2.2 mM, and cells were allowed to grow at 37 °C for an additional 24 h before collection. Cell pellets were harvested by centrifugation, washed in ice-cold phosphate-buffered saline solution and flash-frozen in liquid nitrogen. For cryo-EM analysis, SLC33A1^MAP^ was solubilized in 2% lauryl maltose neopentyl glycol (Anatrace, NG310), 0.2% cholesteryl hemisuccinate Tris salt (Anatrace, CH210), 50 mM HEPES (pH-adjusted with KOH, pH 7.4), 150 mM KCl supplemented with protease-inhibitor cocktail (1 mM phenylmethylsulfonyl fluoride (PMSF), 2.5 mg ml^−1^ aprotinin, 2.5 mg ml^−1^ leupeptin, 1 mg ml^−1^ pepstatin A) and DNase I. For proteoliposome reconstitution, SLC33A1 was solubilized in 2% *n*-dodecyl-β-d-maltoside (DDM; Anatrace, D310), 50 mM HEPES (pH-adjusted with KOH, pH 7.5), 150 mM KCl supplemented with protease-inhibitor cocktail (1 mM PMSF, 2.5 mg ml^−1^ aprotinin, 2.5 mg ml^−1^ leupeptin, 1 mg ml^−1^ pepstatin A) and DNase I. Solubilized proteins were separated by centrifugation 100,000*g* for 1 hr, followed by binding to anti-GFP nanobody resin for 3 h. Anti-GFP nanobody affinity chromatography was carried out by ten column volumes of washing with size-exclusion chromatography (SEC) buffer followed by overnight PreScission digestion and elution with SEC buffer. SEC buffer for cryo-EM analysis consisted of 0.01% lauryl maltose neopentyl glycol, 18-36 μM GDN (anatrace GDN101), 0.001% cholesteryl hemisuccinate Tris salt, 50 mM HEPES (pH-adjusted with KOH, pH 7.5) and 150 mM KCl. The SEC buffer for proteoliposome reconstitution consisted of 0.34 mM DDM, 50 mM HEPES (pH-adjusted with KOH, pH 7.5) and 150 mM KCl. Eluted proteins were concentrated to a volume of 250 μl using Amicon® Ultra-15 Centrifugal Filter Unit (30-kDa cutoff; UFC903008), followed by centrifugation at 21,130*g* for 15 min. Concentrated protein was further purified by SEC on a Superose® 6 Increase 10/300 GL (Cytiva 29-0915-96, 10/300 GL) in SEC buffer. For proteoliposome reconstitution, the peak fractions were pooled and used for proteoliposome reconstitution immediately. For cryo-EM analysis, peak fractions were pooled, mixed and concentrated to protein concentration of 10 mg ml^−1^. Concentrated fractions were then incubated with 20mM GSSG (Thermo J63715.06) and PMab1 Fv-clasp at a 1:1.5 molar ratio for 30 min at room temperature.

### PMab1 Fv-clasp Production and Purification

Sequences for PMab-1 in the Fv-clasp(v2) format were obtained as previously published^32^. The heavy and light chains were cloned in a pETDuet vector (Addgene plasmid #136144) with a N terminus pelB signal sequence for both the heavy and light chain. The heavy chain has a C-terminus 6x his tag.

Expression of Fv-clasp was carried out in BL21-Codon Plus (DE3) E. coli cells (Thermo EC0114) harboring the pETDuet Pmab1 Fv-clasp vectors described above. A single colony was selected and cultured overnight at 37 °C in 10 mL of terrific broth (TB) containing 100 μg/mL ampicillin. The 10 mL overnight E. coli culture was used to inoculate a 1L culture of TB containing 100 μg/mL ampicillin. Cells were cultured at 37 °C until OD600 = 0.7 – 0.9. Protein expression was induced by addition of 0.1 mM IPTG and cultured for an additional 16 – 18 hrs at 25 °C. Cells were harvested by centrifugation (7650 g for 30 min) and the periplasmic E. coli fraction was extracted via osmotic shock as previously described^60,61^. Briefly, harvested cells were suspended in a hypertonic solution of 30 mM Tris, 20% w/v sucrose, 1 mM EDTA, pH 8 (25 mL) and incubated for 30 min at 4 °C. Cells were centrifuged and the supernatant collected. Cells were resuspended in a hypotonic solution of 5 mM MgSO4 (25mL) and incubated for 30 min at 4 °C followed by an additional centrifugation. The supernatant from the hypotonic solution was combined with the supernatant from the hypertonic solution, centrifuged to remove debris, and dialyzed against 20 mM Tris, pH 7.5 and 150 mM NaCl (Tris-buffered saline, TBS) overnight at 4 °C.

The periplasmic solution containing soluble Pmab1 Fv-clasps was clarified over a 0.45 μm filter and purified by Ni^2+^ affinity chromatography as follows. Pmab1 Fv-clasps purified using a two-step chromatography approach. First, the periplasmic fractions were bound to HisPur™ Ni-NTA Resin (Thermo 88222) and then washed sequentially with TBS (20 mM Tris, 150 mM NaCl) containing 20 mM and 40 mM imidazole, followed by elution with TBS containing 500 mM imidazole. Eluted proteins were concentrated using an Amicon® Ultra-15 Centrifugal Filter Unit (10 kDa cutoff) and further purified by size exclusion chromatography on a Superdex 200 Increase 10/300 gel filtration column (Cytiva 28990944) in TBS buffer.

### Electron microscopy sample preparation and data acquisition

A 3 μl volume of 5 mg ml^−1^ purified SLC33A1^MAP^ was mixed with Pmab1 Fv-clasp at a 1:1.5 molar ratio and 20mM GSSG, incubated for 30 min on ice and then applied to glow-discharged Au 400 mesh QUANTIFOIL R1.2/1.3 holey carbon grids (Quantifoil) or Graphene Oxide Au 400 mesh QUANTIFOIL R1.2/1.3 grids before being plunged into liquid-nitrogen-cooled liquid ethane using an FEI Vitrobot Mark IV (FEI Thermo Fisher). For samples frozen on holey carbon grids, the Vitrobot was maintained at 4 °C with 100% humidity, using blotting times of 3.5 s to 4.5 s, blotting force of 0, and a waiting time of 10 s. For samples frozen on graphene oxide grids, the SLC33A1^MAP^ Fv-clasp mixture was diluted 100-fold to a final concentration of 0.05 mg ml^−1^ and the vitrobot was maintained at 4 °C with 100% humidity, using blotting times of 3.5 s to 4.5 s, blotting force of 0, and a waiting time of 35 s. Grids were transferred to a Krios G4 microscope operated at 300 kV and equipped with a Falcon 4i direct electron detector and Selectris X energy filter (Thermo Fisher Scientific).

### Electron microscopy data processing

Images were recorded for 2.93 s using EPU with a total dose of 60 e/Å^2^ with a pixel size of 0.725 Å. A total of three datasets comprising 9,799, 7,072, and 13,336 micrographs were recorded. The images were fractionated into 45 frames, gain-corrected, and aligned using whole-frame and local-motion-correction algorithms by cryoSPARC (v4.5.0)^62,63^. Particles were initially selected using blob picking in cryoSPARC (v4.5.0)^62^. False-positive selections were removed by 2D classification, which allowed generation of an initial model using the *ab initio* algorithm and the training of a Topaz model in cryoSPARC (v4.5.0)^64^. Topaz trained-based autopicking in cryoSPARC (v4.5.0) was implemented to select particle images, resulting in 3,216,504, 3,182,579, and 6,617,058 particles respectively from each dataset^62,64^. False-positive selections and contaminants were excluded through iterative rounds of heterogeneous classification using three references and four decoy classes generated from noise particles through *ab initio* reconstruction in cryoSPARC (v4.5.0), resulting in stacks of 877,231, 815,775, and 906,048 particles from each dataset. At this point these particles from all three of the datasets were combined. Further iterative rounds of heterogeneous classification were used to remove false-positive selections and contaminants yielding 597,738 particles. The selected particles were then subjected to reference-based motion correction in cryoSPARC (v4.5.0), followed by several rounds of heterogeneous refinement using volumes of different states of SLC33A1, resulting in a final stack of 215,071 particles. A consensus reconstruction at a resolution of 3.23 Å was calculated using blush regularization in Relion^65^. After Blush regularization, a focus mask that covers the SLC33A1^MAP^ density but excludes the detergent micelle density, which we refer to as an SLC33A1^MAP^ TMD focus mask was used to for local refinement of the transmembrane domain, yielding a construction at a resolution of 3.26 Å.

### Model building and coordinate refinement

An initial structure of human SLC33A1 was generated with AlphaFold2^66^. The model was docked into the SLC33A1^MAP^ TMD focus refined map using ChimeraX^67^. The model was manually rebuilt to fit the density and GSSG was added in coot^68^. Atomic coordinates were refined against the density-modified map using phenix.real_space_refinement with geometric and Ramachandran restraints maintained throughout^68^. The final model contains residues 70-282 and 293-550 and one bound GSSG molecule.

### Proteoliposome reconstitution

Liposomes were prepared as previously described^69^. Briefly, 1-palmitoyl-2-oleoyl-glycero-3-phosphocholine (PC, Avanti Polar Lipids, 850457), 1-palmitoyl-2-oleoyl-sn-glycero-3-phosphoethanolamine (PE, Avanti Polar Lipids, 850757), and liver L-α-phosphatidylinositol (Liver PI, Avanti Polar Lipids, 840042) in chloroform were mixed at a 5:2:1 (w/w) ratio for Figure 3d and Extended Data Figure 4l, or PC:PE were mixed at a 5:2 (w/w) ratio for Extended Data Figure 7f. Lipids were dried under a rotary evaporator to form a thin film. The lipids were left in a dry-seal vacuum desiccator connected to vacuum overnight. Next, the lipids were rehydrated in buffer containing 150mM KCl, 50 mM HEPES buffer (pH 7.4) at a concentration of 8 mg/ml by shaking until fully dissolved, followed by 6 freeze-thaw cycles. The liposomes were extruded 11 times through 400-nm polycarbonate membranes (Avanti Polar Lipids) using a syringe extruder (Avanti Polar Lipids) to form unilamellar liposomes. Liposomes were destabilized with Triton X-100 at 1:2 (w:w) detergent to lipid ratio and then incubated with purified SLC33A1 at a 1:500 (w/w) protein to lipid ratio for 30 min at room temperature. For the protein-free liposome reconstitution, an equivalent volume of SEC buffer from the protein purification step was added. Detergent was removed by adding Bio-Beads SM-2 (100 mg/ml) (Bio-Rad, 1523920) and rotating for 2 hours at room temperature followed by five additional rounds at 4°C every two hours. The proteoliposome aliquots of 100 µl at a concentration of 8 mg/ml were flash-frozen with liquid nitrogen and stored at −80 °C until use.

### Proteoliposome validation

To ensure efficient protein reconstitution, 100 µl frozen proteoliposomes were thawed and a 10 µl aliquot was collected for Western blot analysis. The remaining sample was centrifuged at 125,000g for 1 hr, and 10 µl of the resulting supernatant was collected for Western blot and the remaining was carefully aspirated. The pellet was resuspended in liposome solution buffer (50 mM HEPES pH 7.4 and 150 mM KCl) containing 2% (w/v) DDM to a final volume of 100 µl. Proteoliposomes were solubilized at 4°C for 1 hr with mild rotation, and a 10 µL sample was collected for Western blot. The remaining solution was centrifuged at 125,000g for 1 hr. 10 µl of the resulting supernatant was collected for Western blot and the remaining was carefully aspirated. The pellet was resuspended in 100 µl of 1× SDS sample buffer. All collected samples were diluted to a protein concentration of 1 ng/µl, and 5 µl of each sample was loaded on a Tris-Glycine gel (Novex™ Tris-Glycine Mini Protein Gels, 10–20%, 1.0 mm, WedgeWell™) for Western blot analysis.

### Proteoliposome counterflow assays

Frozen proteoliposome and protein-free liposome aliquots (lipid concentration: 8 mg/ml) were thawed and diluted twofold in internal liposome buffer with final concentrations of 150mM KCl, 50mM HEPES (pH 7.4), 5 mM (Figure 3a and Extended Data Figure 4l) or 10 mM (Extended Data Figure 7F) unlabeled GSSG (Thermo J63715.06). The liposomes were then subjected to three freeze–thaw cycles and extruded 21 times through 400-nm polycarbonate membranes (Avanti Polar Lipids, 10417104) using a syringe extruder (Avanti Polar Lipids, 610000). After centrifuged at 125,000 x g for 1 hour, the supernatant was removed, and pellets were diluted to a final lipid concentration of 2.5 mg/ml in wash buffer consisting of 150mM KCl and 50mM HEPES (pH 7.4). For each sample, 25 µl of liposomes were mixed with 25 µl of the 2 x reaction buffer consisting of 10 µM [^3^H]GSSG (Figure 3a and Extended Data Figure 4l) or 40 µM [^3^H]GSSG (Extended Data Figure 7F), 150 mM KCl, and 50 mM HEPES (pH 7.4). For GSSG competition assay (Extended Data Figure 4l), the 2x reaction buffer consisted of 10 μM [^3^H]GSSG, 5mM GSSG, 150mM KCl, and 50mM HEPES (pH 7.4). After incubated for the time points described in the figure legends at room temperature, the reactions were arrested by adding the proteoliposome solution to 200 µl ice-cold wash buffer and aspirating it through 0.22-μm membrane filters (Millipore, MSGVN2210) under vacuum. The membrane filters were washed five times with a 200 µl ice-cold wash buffer and transferred to a scintillation vial with 5 ml of Insta-Gel Plus scintillation cocktail (Perkin Elmer, 601339). Radioactivity was measured with a liquid scintillation analyzer (Perkin Elmer). Proteoliposome counterflow assays were repeated five times. For competition assays, unlabeled GSSG ((0.1 to 10 mM)) or acetyl-CoA (5 µM to 12.5 mM) were added to the external solution.

### Thermal Shift assays

Tryptophan fluorescence measurements were carried out using Prometheus Panta (NanoTemper Technologies). SLC33A1 was diluted in dilution buffer containing 50 mM HEPES pH 7.4, 150 mM NaCl and 0.001% (w/v) LMNG to a concentration of 2.66 μM. 10 μl of SLC33A1 was mixed with equal volumes of two-fold serially diluted solutions of various molecules (6000 μM to 0.732 μM) to achieve a final protein concentration of 1.33 μM of SLC33A1, followed by room temperature incubation for 20 min. 10 μl of protein-compound mixture was used per Prometheus high-sensitivity capillary (NanoTemper Technologies). Recorded tryptophan fluorescence (350 nm/330 nm) was analysed by using NanoTemper instrument software. The first derivative was calculated from each curve to determine the melting temperature. Three technical replicates were recorded for data analysis. Thermal shift measurements were repeated two times.

### Molecular Dynamics simulations

#### Simulation system preparation

All molecular dynamics (MD) simulations were conducted using OpenMM 7.7.0^70^. The cryo-EM structure of GSSG-bound SLC33A1^MAP^with bound GSSG was initially prepared for simulations using the Protein Preparation Wizard at pH 7.5 within Schrödinger Maestro release 2022-1. The position of SLC33A1^MAP^ in a membrane was calculated using the Orientation of Proteins in Membranes (OPM) PPM 2.0 web server^71^. GSSG was modeled as partially protonated with a net charge of –2 and was parameterized with the CHARMM General Force Field (CGenFF)^72^ using the CHARMMGUI^73^. Ligand Reader and Modeler^74^. The resultant GSSG atom naming scheme was used to replace the original GSSG atom names in SLC33A1^MAP^ PDB files to maintain consistency with further CHARMMGUI protocols. The SLC33A1^MAP^-GSSG system was then prepared for simulation using the CHARMMGUI bilayer builder^75,76^. Termini were capped, glu187 was set to the partially protonated state based on protonation predictions within Maestro, and the systems were embedded in POPC: POPE = 3:1 membrane bilayers with equal numbers of lipids on both leaflets. The membrane embedded system was placed in water boxes containing explicit water based on the TIP3P water model with neutralizing 150 mM KCl. Periodic boundary conditions were established with a box size of 125 x 125 x 119 Å^75^. Further simulation inputs for OpenMM were generated.

#### Simulation system equilibration and energy minimization

Following preparation, the systems were energy minimized with an energy tolerance of 100 kj/mol and equilibrated in six sequential steps involving loosening of positional restraints on lipid atoms, protein backbone atoms, and protein side chain atoms (equilibration parameters in Supplementary Table 5). Equilibration was conducted using a Langevin Integrator in a NPT ensemble^77^. Systems at the final step of this equilibration were subsequently used to seed production simulations on the Memorial Sloan Kettering Cancer Center computational cluster, resulting in an aggregate simulation time of 3.4 µs. Production simulations were conducted using a LFMiddle Langevin Integrator in a NPT ensemble (simulation parameters summarized in Supplementary Table 6)^78^.

#### Simulation Trajectory analysis

All simulation trajectory analysis was performed using MDTraj^79^ and NumPy^80^. Prior to any measurement, all protein components in the simulation system were aligned to the first frame and centered. Radii of gyration for GSSG were computed by calculating the root-mean-square-distance of the GSSG atoms from its center of mass. Distances between GSSG and Leu335 were computed as the euclidean distance between the center of mass of GSSG and the gamma carbon of Leu335. To quantify the frequency of interaction between GSSG functional groups and protein residues, the pairwise distances between all GSSG functional group atoms and protein atoms were calculated for each trajectory frame. A protein residue was noted to be interacting with GSSG if any of its atoms were within a 4 Å cutoff of any GSSG functional group atom. Interaction frequency was computed as the number of frames that a particular residue interacted with GSSG divided by total number of frames.

### Figures

Figures were prepared with PyMol 2.5.8 (https://www.pymol.org), ChimeraX 1.7.1^68^, GraphPad Prism 10 (https://www.graphpad.com), Matplotlib^81^, and Seaborn^82^.

### Statistics and reproducibility

Statistical analyses were performed using GraphPad Prism (vX). The statistical tests used, definitions of n, and exact p values are indicated in the figure legends. Data are presented as mean ± s.d. or mean ± s.e.m., as indicated. No statistical methods were used to pre-determine sample sizes, but sample sizes were chosen based on prior experience and are consistent with those generally employed in the field. Data distribution was assumed to be normal but this was not formally tested. The experiments were not randomized; samples were assigned to experimental groups based on experimental design and treatment conditions. Data collection and analysis were not performed blind to the conditions of the experiments. For CRISPR/Cas9 genetic screens, gene scores derived from three or fewer guide RNAs were excluded from the analysis, as these provide insufficient statistical confidence for gene-level scoring. For oxidized cysteine reactivity proteomic data, one sample in the Slc33a1_KO + Vector group was excluded due to poor TMT labeling efficiency relative to all other samples. For cryo-EM analysis, micrographs showing high drift or poor CTF fits were excluded during preprocessing, and individual particles were excluded by 2D and 3D classification following standard procedures in single-particle cryo-EM analysis; the particle selection workflow is shown in Extended Data Fig. 6. For all other experiments, no data were excluded. All experiments were performed with at least three independent biological replicates unless otherwise stated in the figure legends.

## Data Availability

ER-IP RNA–seq data and KPK RNA-Seq have been deposited in the Gene Expression Omnibus (GEO) under accession codes GSE318826 and GSE318827. Proteomics data have been deposited in PRIDE under accession number PXD074236 and PXD074465. Genetic screen results are shared in Supplementary Table 7-8. Metabolomics and lipidomics datasets supporting the findings of this study are provided in Supplementary Table 12. Cryo-EM maps have been deposited in the Electron Microscopy Data Bank (EMDB) under the accession codes EMDB-48699 (composite map) and EMDB-48638 (local refinement map). Atomic coordinates have been deposited in the Protein Data Bank (PDB) under the accession codes 9MUN (GSSG-bound SLC33A1^MAP^). Source data have been provided in Source Data. All other data supporting the findings of this study are available from the corresponding author on reasonable request.

## Code Availability

All atomic models and code associated with molecular dynamics simulations can be found at https://github.com/choderalab/slc33a1_md.

---

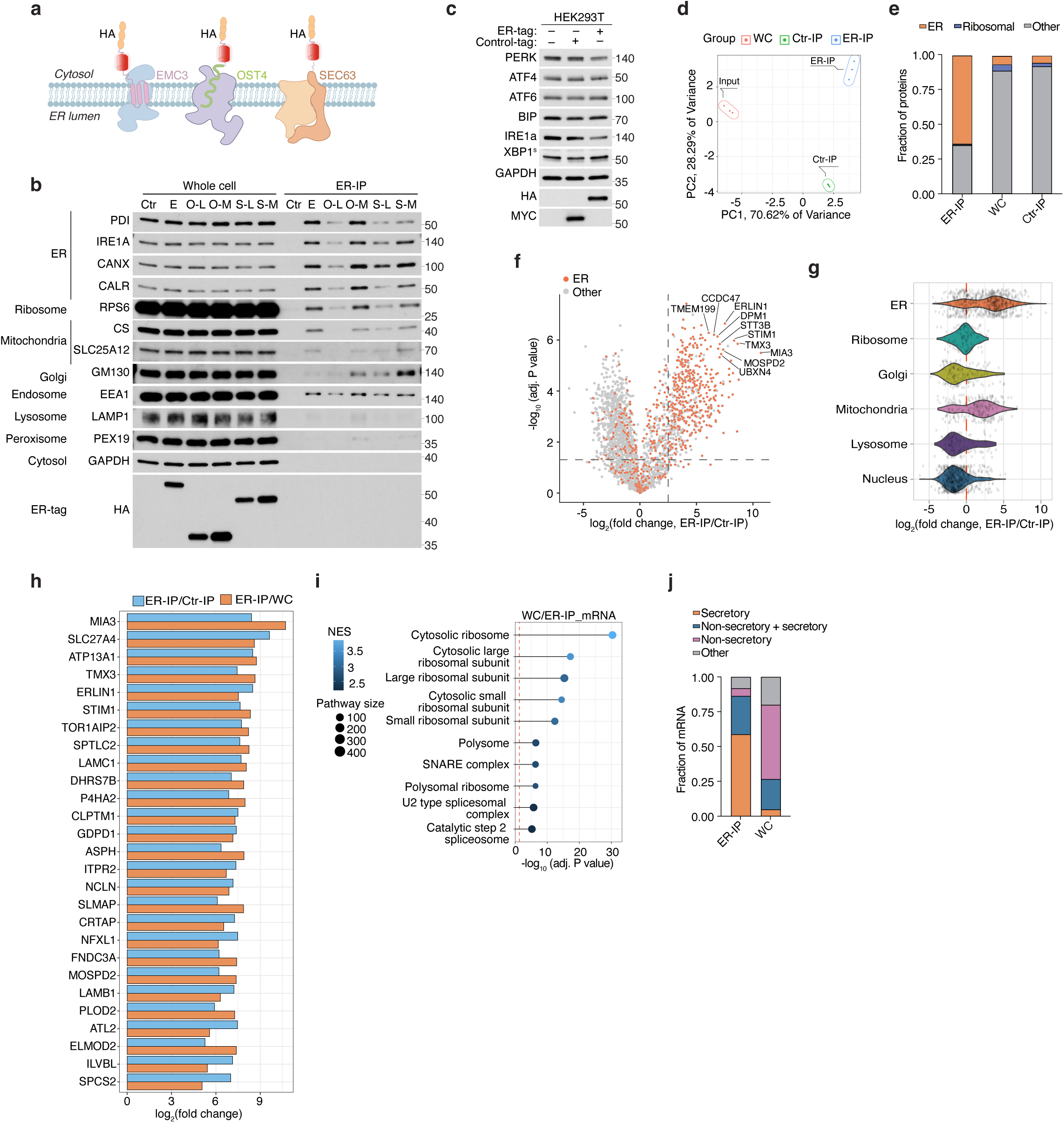

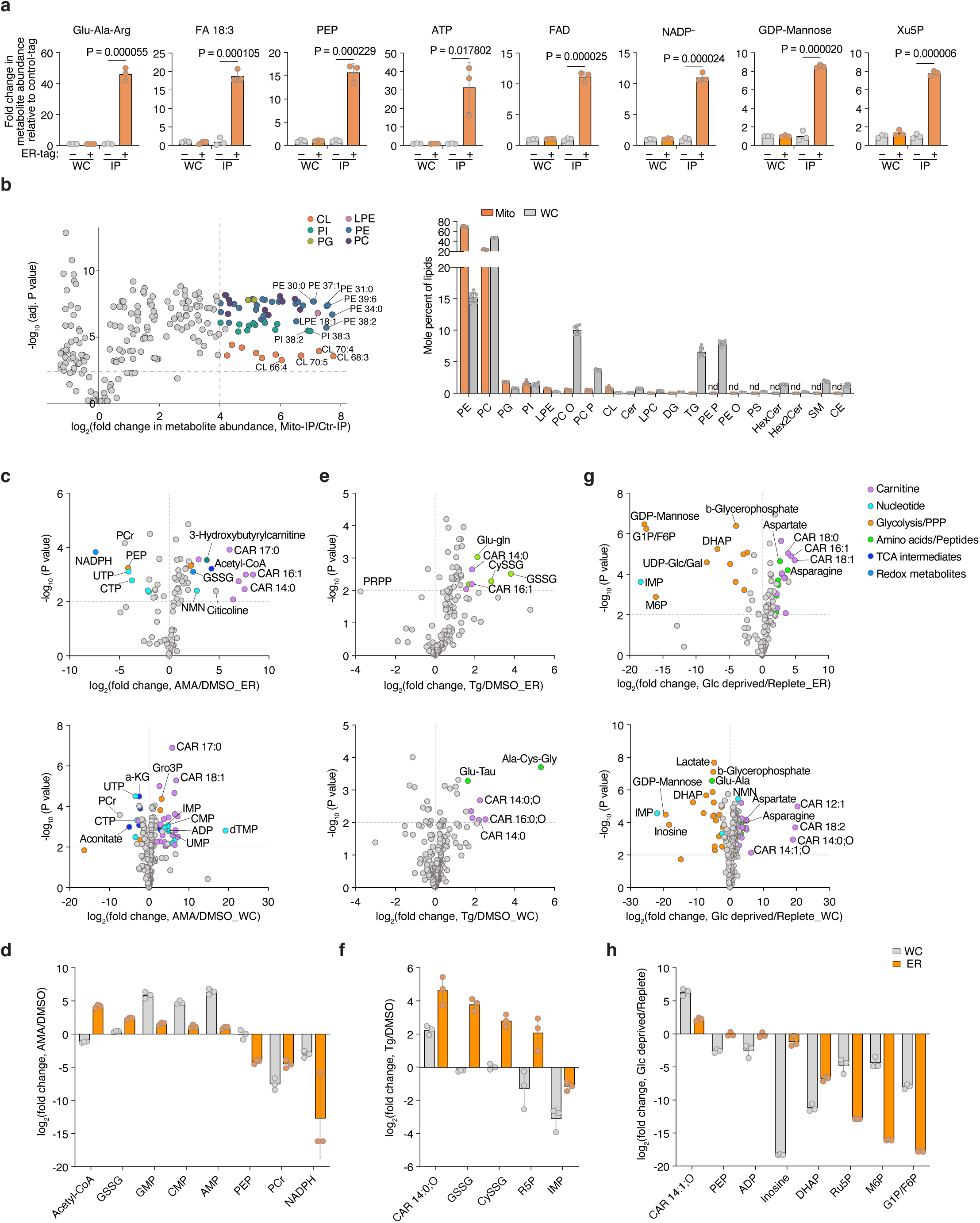

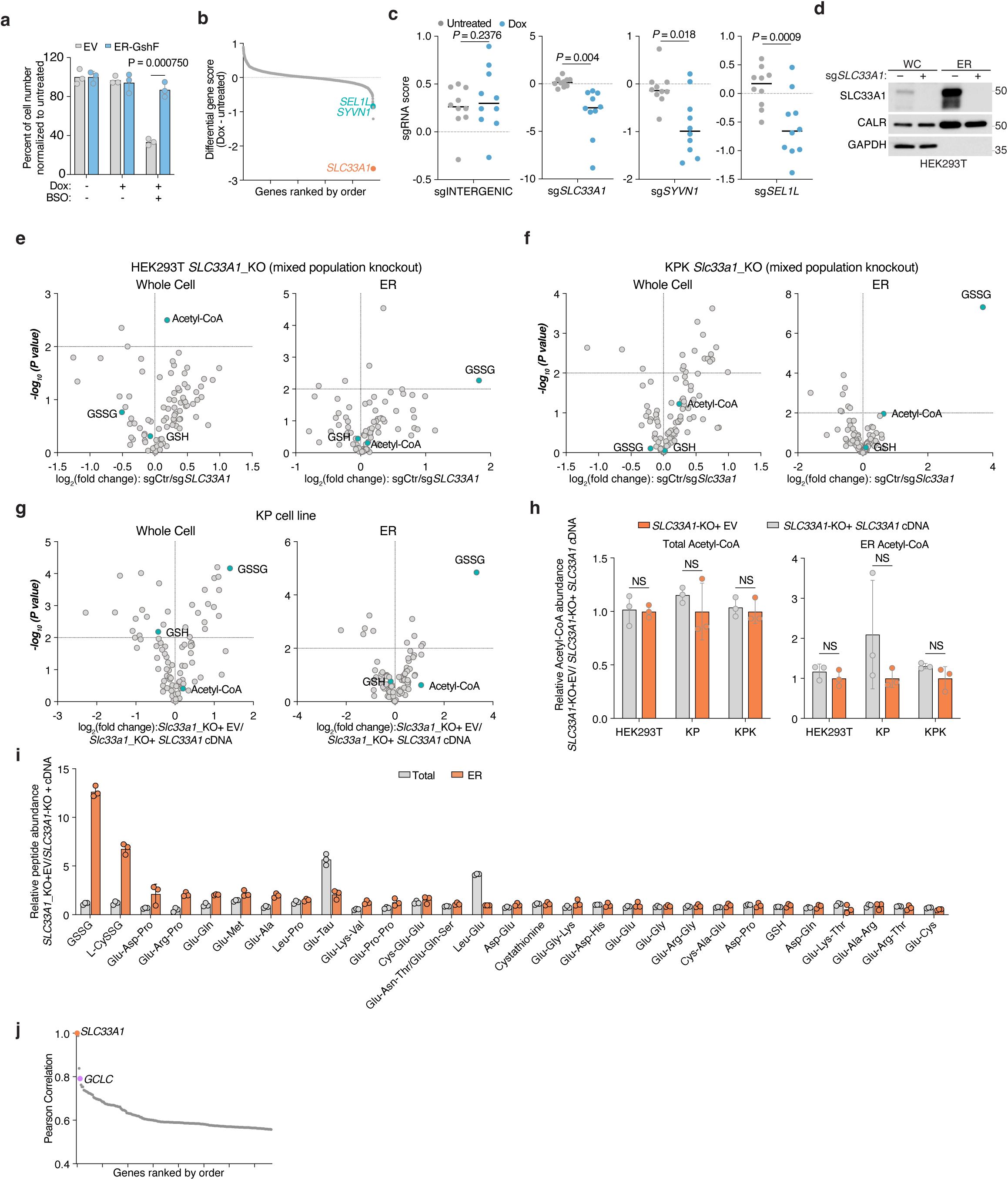

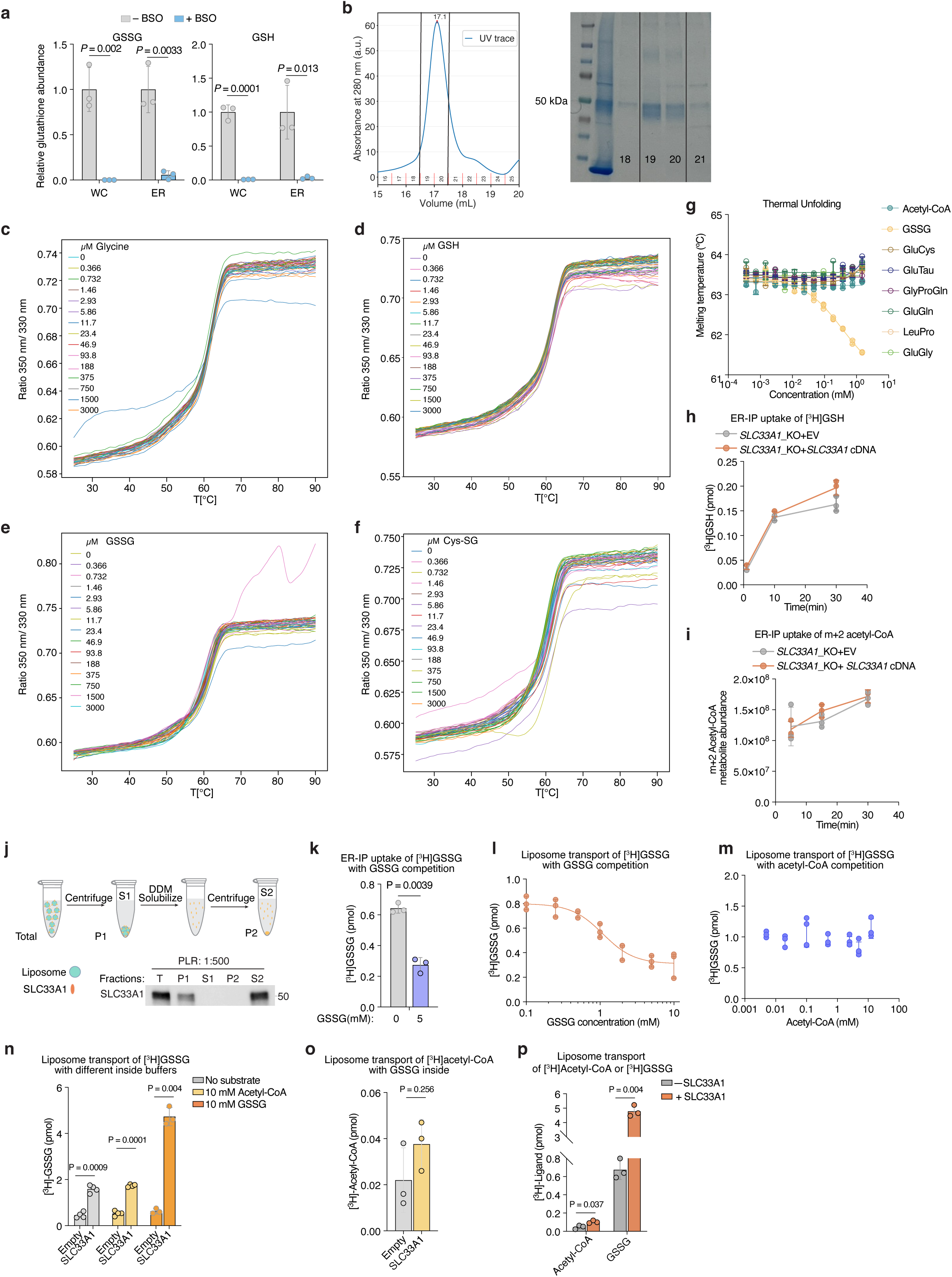

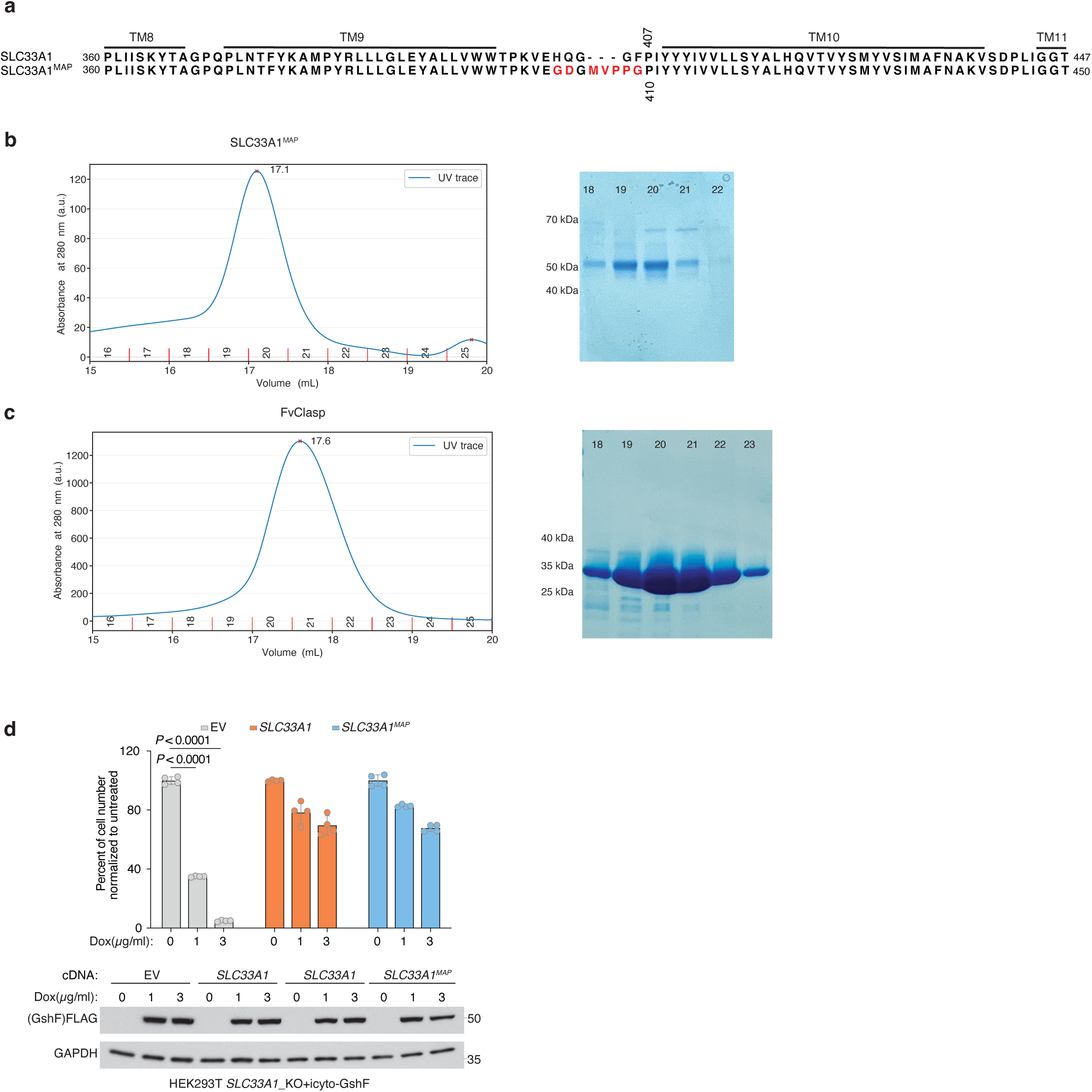

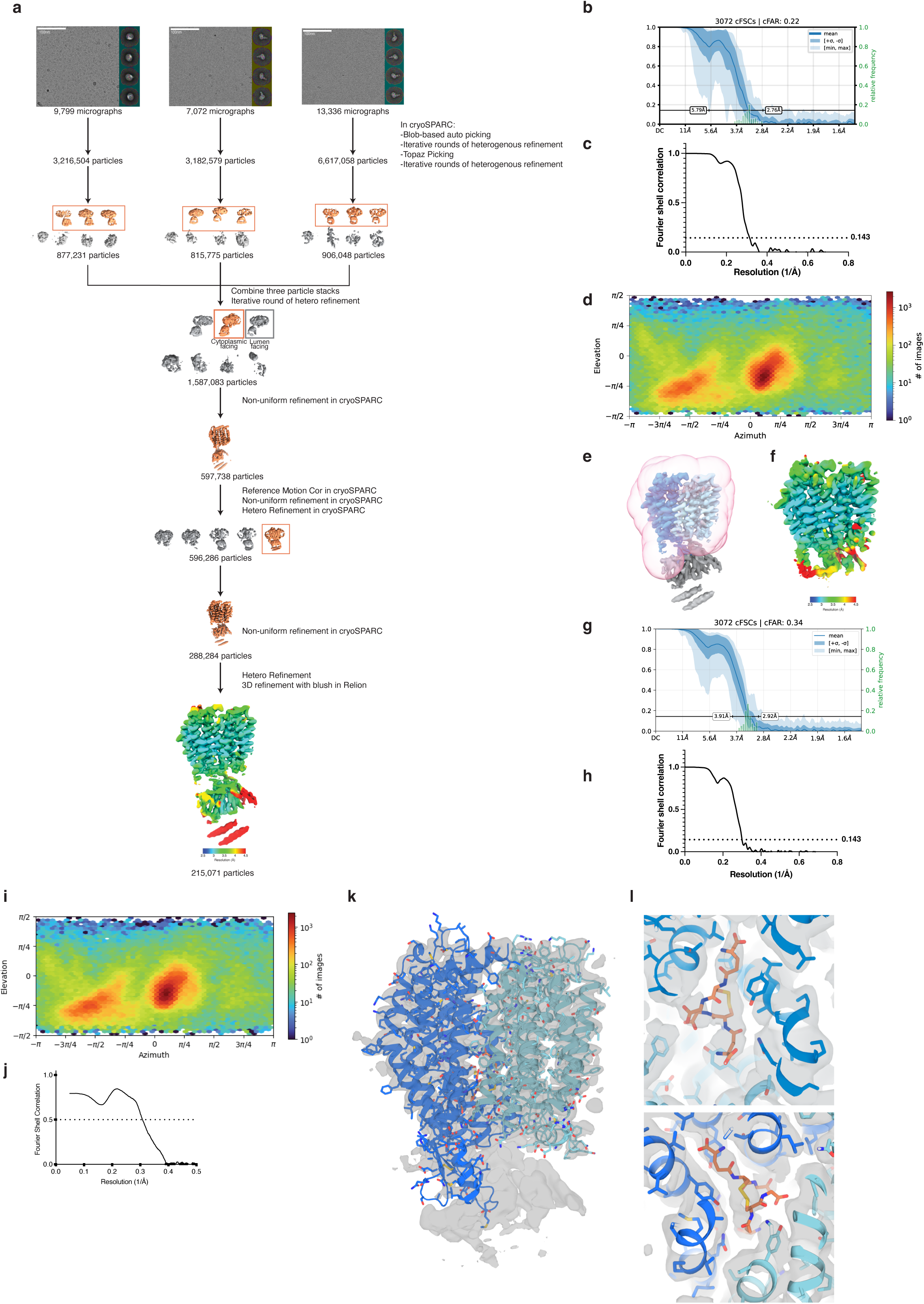

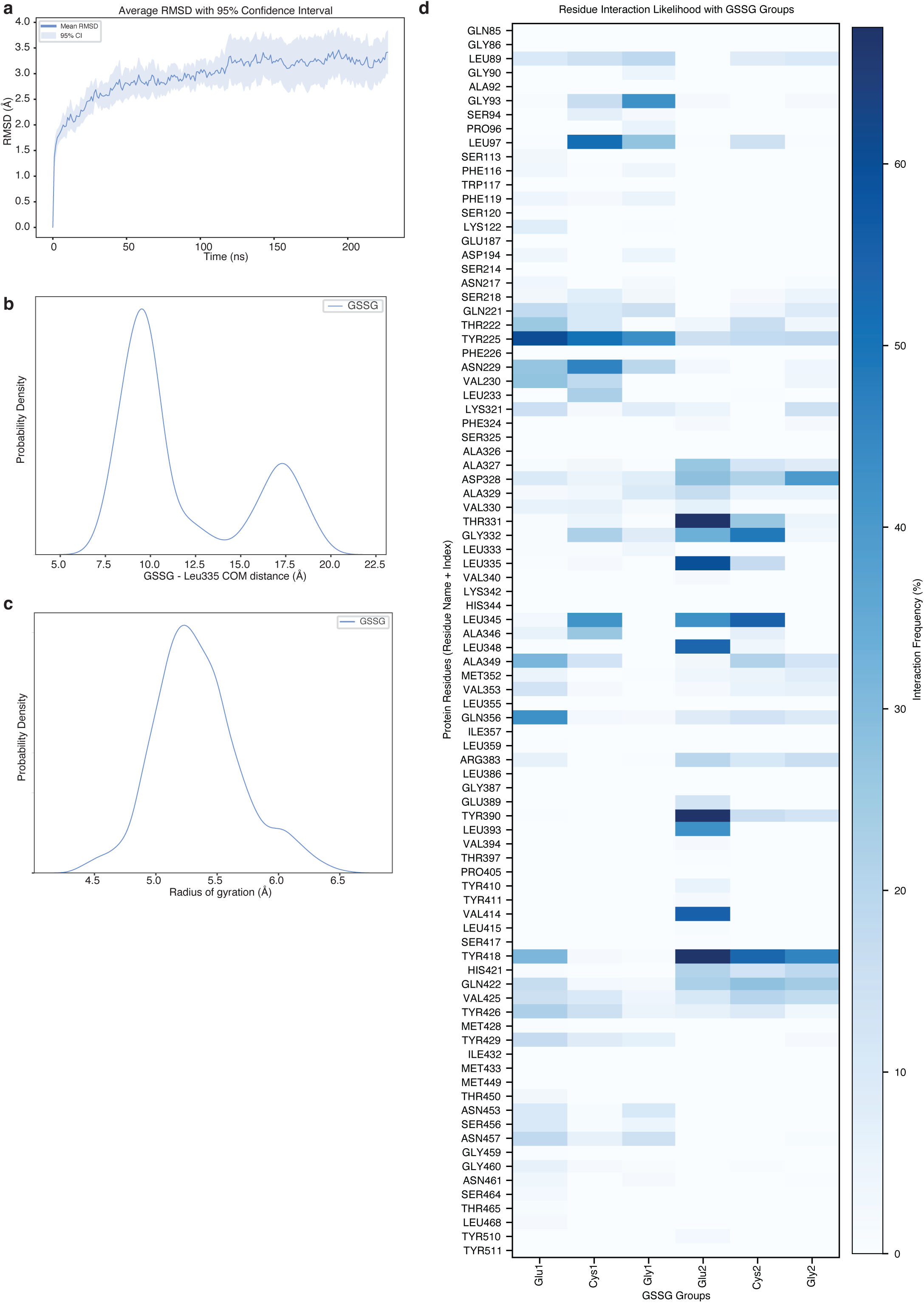

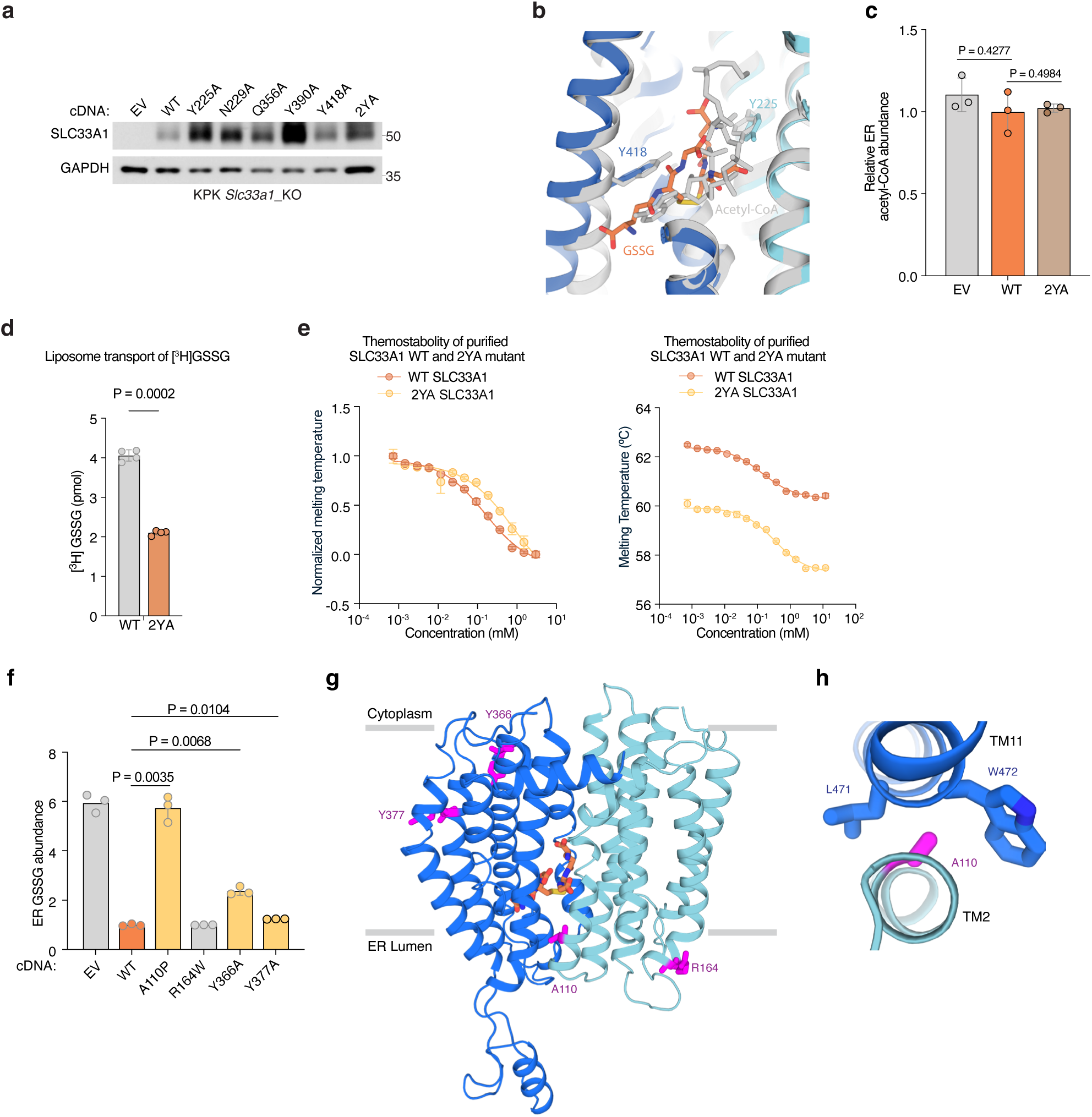

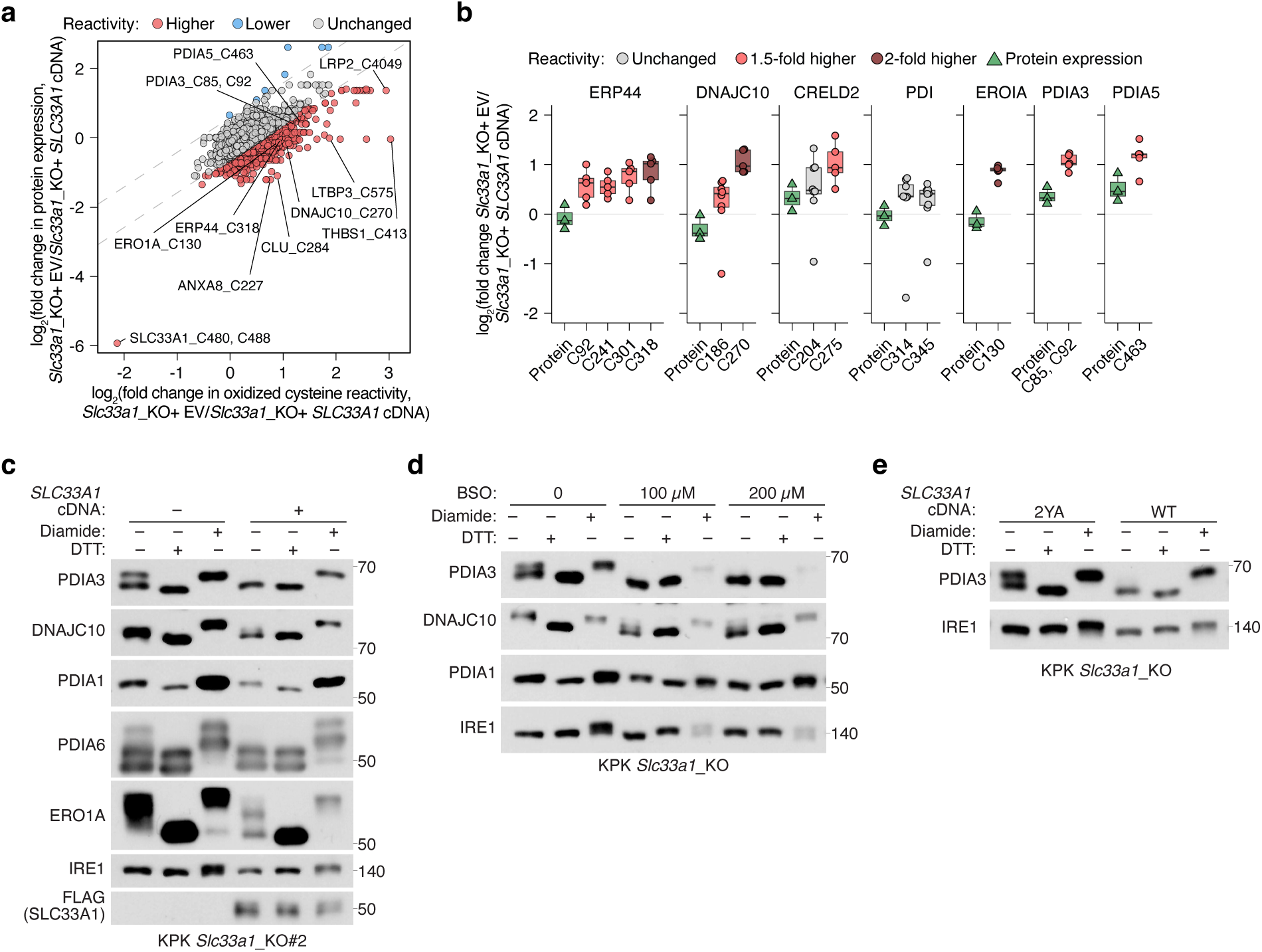

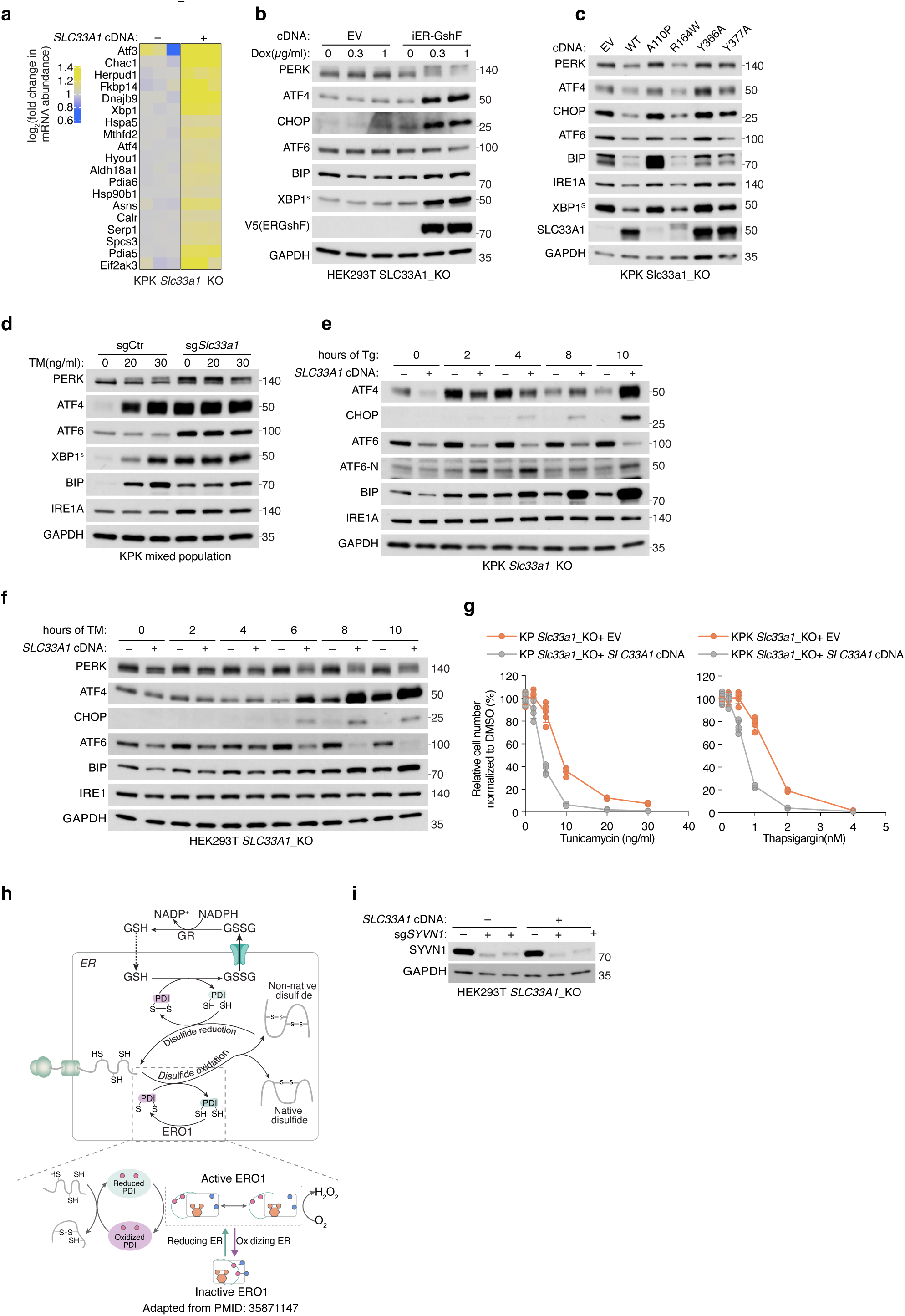

